# A Bayesian framework for unifying data cleaning, source separation and imaging of electroencephalographic signals

**DOI:** 10.1101/559450

**Authors:** Alejandro Ojeda, Marius Klug, Kenneth Kreutz-Delgado, Klaus Gramann, Jyoti Mishra

## Abstract

Electroencephalographic (EEG) source imaging depends upon sophisticated signal processing algorithms to deal with the problems of data cleaning, source separation, and localization. Typically, these problems are sequentially addressed by independent heuristics, limiting the use of EEG images on a variety of applications. Here, we propose a unifying empirical Bayes framework in which these dissimilar problems can be solved using a single algorithm. We use spatial sparsity constraints to adaptively segregate brain sources into maximally independent components with known anatomical support, while minimally overlapping artifactual activity. The framework yields a recursive inverse spatiotemporal filter that can be used for offline and online applications. We call this filter Recursive Sparse Bayesian Learning (RSBL). Of theoretical relevance, we demonstrate the connections between Infomax Independent Component Analysis and RSBL. We use simulations to show that RSBL can separate and localize cortical and artifact components that overlap in space and time from noisy data. On real data, we use RSBL to analyze single-trial error-related potentials, finding sources in the cingulate gyrus. We further benchmark our algorithm on two unrelated EEG studies showing that: 1) it outperforms Infomax for source separation on short time-scales and 2), unlike the popular Artifact Subspace Removal algorithm, it can reduce artifacts without significantly distorting clean epochs. Finally, we analyze mobile brain/body imaging data to characterize the brain dynamics supporting heading computation during full-body rotations, replicating the main findings of previous experimental literature.

## 1. Introduction

The electroencephalogram (EEG) is a noninvasive functional brain imaging modality that allows the study of brain electrical activity with excellent temporal resolution. Compared to other noninvasive imaging modalities such as fMRI, PET, SPECT, and MEG, EEG acquisition can be mobile and more affordable (Mcdowell et al., 2013; Mehta and Parasuraman, 2013), allowing the widespread study of human cognition and behavior under more ecologically valid experimental conditions (Makeig et al., 2009). Imaging cognitive processes while participants engage naturally with their environment (natural cognition in action (Gramann et al., 2014)) has potential for developing a new generation of applications in brain-computer interfaces (BCI), mental health, rehabilitation, and neuroergonomics (Mishra and Gazzaley, 2014; Mishra et al., 2016; Jungnickel and Gramann, 2016; Wagner et al., 2016). However, despite impressive methodological advances in the estimation of the electrical activity of the cortex from EEG voltages recorded on the scalp, a number of practical and theoretical issues remain unsolved.

Imaging EEG source activity (also known as electromagnetic source imaging or ESI) is challenging for several reasons. First, since many configurations of currents in the brain can elicit the same EEG scalp topography (Michel and Murray, 2012), it entails solving an ill-posed inverse problem (Lopes da Silva, 2013). Second, the EEG signal is often contaminated by artifacts of non-brain origin such as electrooculographic (EOG) and electromyographic (EMG) activity that need to be identified and removed. Third, due to the low spatial resolution of the EEG, traditional inverse solvers produce estimates that can be a (distorted) mixture of the true source maps (Biscay et al., 2018). These problems are usually addressed separately using a variety of heuristics, making it difficult to systematize a methodology for obtaining biologically plausible single-trial EEG source estimates in the presence of artifacts. The objective of this paper is to develop a unifying Bayesian framework in which these, apparently dissimilar, problems can be understood and solved in a principled manner using a single algorithm.

The problem of EEG source estimation is even harder if we consider that there is evidence that brain responses are generated by time-varying network dynamics that can exhibit nonlinear features (Breakspear, 2017; Khambhati et al., 2018), which renders the simplifying assumptions of linearity and stationarity used by most inverse methods hard to justify. Thus, the objective of our framework is to produce a spatiotemporal inverse filter that can map each EEG sample to the source space, minimizing source mixing, and factoring out the corrupting effect of artifacts in an adaptive manner.

To cope with the ill-posed nature of the inverse problem and ensure functional images with biological relevance, several inverse algorithms have been proposed that seek to estimate EEG sources subject to neurophysiologically reasonable spatial (Haufe et al., 2011; Friston et al., 2008; Trujillo-Barreto et al., 2004; Pascual-Marqui et al., 2002; Baillet et al., 2001; Hämäläinen and Ilmoniemi, 1994), spatiotemporal (Martínez-Vargas et al., 2015; Valdés-Sosa et al., 2009; Trujillo-Barreto et al., 2008), and frequency-domain (Gramfort et al., 2013) constraints, just to mention a few examples. These approaches can work relatively well when the EEG samples are corrupted by Gaussian noise and the signal to noise ratio (SNR) is high. In practice, however, raw EEG data are affected by many other types of noise such as interference from the 50/60 Hz AC line, pseudo-random muscle activity, and mechanically induced artifacts, among others. Thus, before source estimation, non-Gaussian artifacts need to be removed from the data.

There is a plethora of methods for dealing with artifacts corrupting the EEG signal (see reviews by Mannan et al. (2018); Islam et al. (2016)). Popular approaches used in BCI applications are based on adaptive noise cancellation (Kilicarslan et al., 2016) or Artifact Subspace Removal (ASR) (Mullen et al., 2015) algorithms. The former has the inconvenience that an additional channel recording purely artifactual activity (i.e., EOG or EMG activity not admixed with EEG) needs to be provided, while the latter rests on the assumption that the statistics of data and artifacts stay the same after an initial calibration phase. In studies where the data can be analyzed offline, artifactual components can be largely removed using Independent Component Analysis (ICA) (Jung et al., 2000). ICA-based cleaning, however, has the drawback that non-brain components need to be identified for removal, which is usually done manually based on the practitioner’s experience.

ICA is a special case of blind source separation (BSS) method (Cichocki and Amari, 2002) that can be used to linearly decompose EEG data into components that are maximally statistically independent. ICA has been used to analyze event-related potentials (ERP) under the assumptions that during the task 1) the decomposition is stationary and 2) that brain components can be modeled as a predefined number of dipolar point processes with fixed spatial location and orientation (Makeig and Onton, 2011). The stationarity assumption can be relaxed using a mixture of ICA models (Palmer et al., 2011) while the selection of brain scalp projections is typically done either manually or automatically based on the residual variance afforded by a dipole fitting algorithm. The practical use of ICA has been limited by its computational cost and the need for user intervention. Only recently, a real-time recursive ICA algorithm has been proposed (Hsu et al., 2016), as well as a number of automatic methods for minimizing the subjectivity of manual component selection (Tamburro et al., 2018; Pion-Tonachini et al., 2017; Radüntz et al., 2017). Despite these advances, turning ICA into a brain imaging modality requires that after source separation, we solve the inverse problem of localizing the set of identified brain components into the cortical space.

One way of estimating EEG sources subject to multiple assumptions (constraints) in a principled manner is to use the framework of parametric empirical Bayes (PEB) (Morris, 1983; Casella, 1985). In this framework, constraints are used to furnish prior probability density functions (pdfs). Empirical Bayes methods use data to infer the parameters controlling the priors (hyperparameters), such that those assumptions that are not supported by the data can be automatically discarded without user intervention. Here we use priors to “encourage” source images to belong to a functional space with biological relevance, but the exact form of those priors is determined by the data (empirically).

In the context of sparsity-inducing priors, PEB is sometimes referred to as Sparse Bayesian Learning (SBL) (Tipping, 2001). The PEB/SBL framework for EEG/MEG has been proposed for recovering instantaneous responses in the context of event-related potential (ERP) experiments (Owen et al., 2012; Henson et al., 2011; Friston et al., 2008). However, there are many applications of interest where source mapping needs to operate as a filter on the continuous EEG signal (e.g., brain monitoring and BCI). One way of extending the instantaneous approach is to introduce additional temporal constraints on the source dynamics in the form of a spatiotemporal prior. With the inclusion of a source dynamics model, the probabilistic generative model (PGM) of the EEG signal can be naturally expressed in the state-space framework (Kalman, 1960).

In recent decades, the state-space framework has been exploited by several authors to solve the inverse problem of the EEG in online fashion. Yamashita et al. (2004) proposed modeling the source dynamics with a nearest neighbor autoregressive model, leading to a Recursive Penalized Least Squares (RPLS) algorithm. Galka et al. (2004) extended RPLS by adding a spatial whitening transformation that made the application of the Kalman filter tractable. Lamus et al. (2012) used a source model similar to the one in (Yamashita et al., 2004) to implement a Kalman filter with online hyperparameter updates. Such an approach, however, can be computationally intractable due to the need to compute a high-dimensional source (state) covariance matrix. Long et al. (2011) sidestepped this limitation with an implementation that can take advantage of a supercomputer parallel architecture. In the presence of oversimplified neurodynamical models, however, the use of the standard Kalman filter formulas may lead to accumulation of errors and divergent behavior (Anderson and Moore, 2012). We expand on this point in Section 2.4.

In this paper, we extend the pioneering work of Yamashita et al. (2004) along two important directions. First, we augment the PGM of the EEG to include the effects of non-brain (artifact) source dynamics. Second, we use spatial sparsity constraints to adaptively segregate brain sources into maximally independent components while minimally overlapping artifactual activity. Henceforth, we refer to this new approach as Recursive Sparse Bayesian Learning (RSBL). Our main contribution is that, by explicitly modeling non-brain sources, we can unify three of the most common problems in EEG analysis: data cleaning, source separation, and source imaging. Furthermore, we show that by updating the parameters of our model online, we can adapt the spatial resolution of our inverse filter so that each EEG sample is optimally localized without temporal discontinuities, thereby showing potential for tracking non-stationary brain dynamics. On the theoretical side, we point out the connections between distributed source imaging and ICA, two popular approaches that are often perceived to be at odds with one another.

Throughout this paper we use lowercase and uppercase bold characters and symbols to denote column-vectors and matrices respectively, **â** is an estimate of the parameter vector **a**, and **I**_*N*_ is a *N* × *N* identity matrix.

## 2. Methods

It has been shown that popular instantaneous source estimation algorithms used in ESI such as weighted minimum *l*_2_-norm (Baillet et al., 2001), FOCUSS (Cotter et al., 2005; Gorodnitsky and Rao, 1997), minimum current estimation (Huang et al., 2006), sLORETA (Pascual-Marqui et al., 2002), beamforming (Van Veen et al., 1997), variational Bayes (Friston et al., 2008), and others can be expressed in a unifying Bayesian framework (Wipf and Nagarajan, 2009). We extend this framework by 1) explicitly modeling non-brain artifact sources and 2) introducing a temporal constraint to yield continuous source time series estimates.

### 2.1. Cortical and artifact source modeling

In source imaging, the neural activity is often referred to as the primary current density (PCD) (Baillet et al., 2001) and it is defined on a dense grid of known cortical locations (the source space). Typically, a vector of *N*_*y*_ EEG measurements at sample 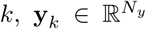, relates to the activity of *N*_*g*_ sources, 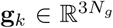, through the following instantaneous linear equation (Dale and Sereno, 1993),

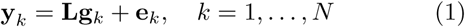

where **g**_*k*_ is the vector of PCD values along the three orthogonal directions and 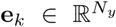 represents the measurement noise vector. The PCD is projected to the sensor space through the lead field matrix 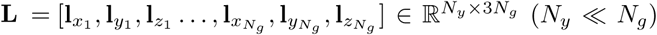 where each column **l**_*i*_ denotes the scalp projection of the *i*th unitary current dipole along a canonical axis. The lead field matrix is usually precomputed for a given electrical model of the head derived from a subject-specific MRI (Hallez et al., 2007). Alternatively, if an individual MRI is not available, an approximated lead field matrix obtained from a high-resolution template can be used (Huang et al., 2016). Then, the instantaneous inverse problem of the EEG can be stated as the estimation of a source configuration **ĝ**_*k*_ that is likely to produce the scalp topography **y**_*k*_.

In the generative model presented above, the noise term **e**_*k*_ is assumed to be Gaussian and spatially uncorrelated with variance *λ*_*k*_. This simplification is acceptable as long as EEG topographies are not affected by non-Gaussian pseudo-random artifacts generated by eye blinks, lateral eye movements, facial and neck muscle activity, body movement, among others. Therefore, before source estimation, EEG data are usually heavily preprocessed and cleaned (Bigdely-Shamlo et al., 2015). Since artifacts contribute linearly to the sensors, ideally, one would like to characterize their scalp projections to describe the signal acquisition more accurately. To this end, we propose the following generalization of Eq (1),

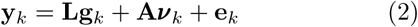

where 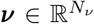 is a vector of *N*_*ν*_ artifact sources and 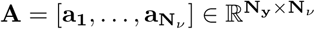 is a dictionary of artifact scalp projections (see Fig 1).

**Figure 1:**
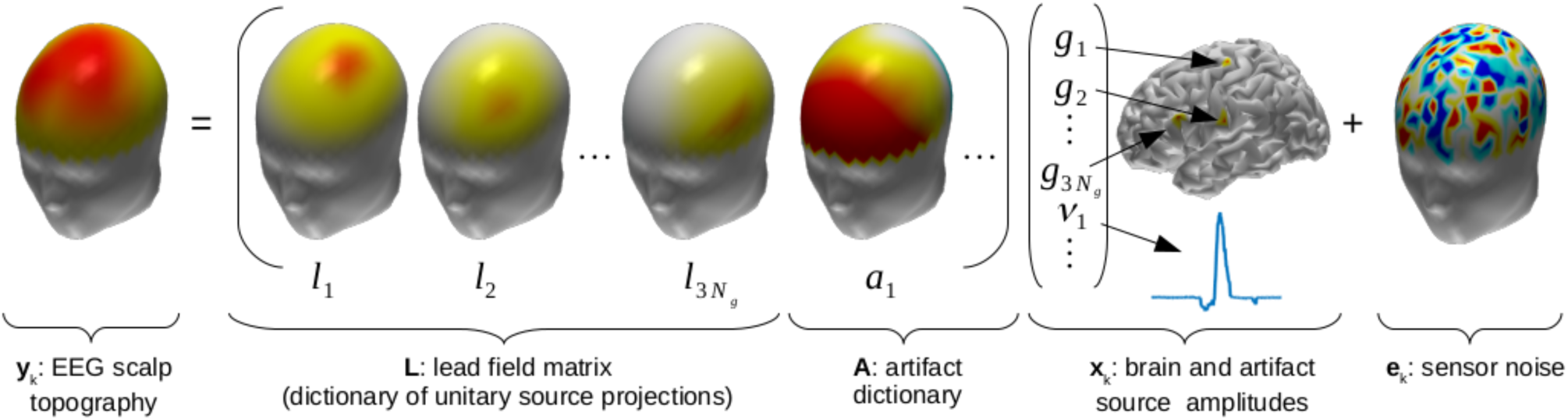
Proposed augmented generative model of the EEG. The model postulates that the EEG scalp topography **y**_**k**_ arises from the linear superposition of brain *g*_*i,k*_ and artifact *ν*_*j,k*_ components weighted by their respective scalp projections **l**_**i**_ and **a**_**j**_, corrupted by spatially uncorrelated Gaussian noise **e**_**k**_.

Although the entries of **A** that correspond to muscle activity may be obtained based on a detailed electromechanical model of the body (Böl et al., 2011), in most studies this approach may not be feasible due to computational and budgetary constraints. Janani et al. (2017) modelled **A** by expanding the lead field matrix to account for the contribution of putative scalp sources, which were assumed to be the generators of EMG activity. They used sLORETA to estimate brain and scalp sources simultaneously. Although this approach was shown to be as effective as ICA-based artifact removal, it was suggested by the authors that the use of the non-sparse solver sLORETA may lead to unrealistic configurations of brain and non-brain sources. Similarly, Fujiwara et al. (2009) augmented the magnetic lead field matrix to model the scalp contribution of two current dipoles located behind the eyes and used a Bayesian approach, that has similarities with ours, to estimate brain and eye source activity from MEG data. Although successful for removing EOG activity, in their formulation, Fujiwara et al. (2009) ignored other types of artifacts that are harder to model such as those produced by muscular activity.

In this paper, we take an empirical view inspired by the success of ICA-based artifact removal approaches. We propose constructing the dictionary **A** using a set of stereotypical artifact scalp projections such as those obtained from running ICA on a database of EEG recordings (Bigdely-Shamlo et al., 2013a). Then we rewrite Eq (2) in a compact manner as follows,

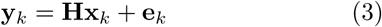

where **H ≜** [**L, A**] is an observation operator and 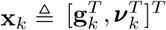 is the augmented vector of hidden (latent) brain and artifact sources (see Fig 1).

Note that, structurally, the standard generative model in Eq (1) and the augmented one in Eq (3) are identical. However, they differ in that in Eq (3) we are explicitly modeling the instantaneous spatial contribution of non-brain sources to the scalp topography **y**_*k*_. Therefore, in theory, we could dispense with computationally expensive preprocessing data cleaning procedures. The assumption of Gaussian measurement noise yields the following likelihood function,

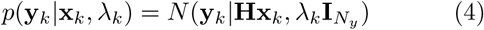

#### 2.1.1. Parameterization of the observation operator

To obtain the observation operator **H** for a given subject, we need head models and channel positions of that subject and those in the EEG database. As we mentioned earlier, a database of stereotypical artifact scalp projections can be obtained by running ICA on a large collection of EEG data sets. The database could include data from different studies, populations, and montages so that a “universal” artifact dictionary can be compiled offline.

Note that the key idea is to use the precomputed artifact dictionary to approximate the scalp projection of stereotypical artifact components of new subjects without actually running ICA on their data. Towards that end, ideally, each subject would have their companion structural MRI and digitized channel locations fully describing the anatomical support on which EEG data were collected. If only sensor positions were available, the procedure proposed by Darvas et al. (2006) could be used to obtain individualized head models using a single template head.

Most EEG studies, however, do not include MRI data or subject-specific sensor positions, but just the sensor labels. Here we propose a co-registration procedure that requires sensor positions only, either measured with a digitizer or pulled from a standard montage file, and a template head model. We use a four-layer (scalp, outer skull, inner skull, and cortex) head model derived from the “Colin27” MRI template with fiducial landmarks (nasion, inion, vertex, left and right preauricular points) and 339 sensors located on the surface of the scalp. The sensors are placed and named according to a superset of the 10/20 system.

Before starting data collection, we use the fiducial points marked on the participant’s head to estimate an affine transformation from the individual space to the space of the template. If fiducial points are not available, a common set of sensors between the template and the individual montage can be used to estimate the transformation. We use the affine transformation to map the montage of the participant onto the skin surface of the template; this is what we call in Fig. 2 “individualized head model”. We compute the lead field matrix **L** using the boundary element method implemented in the software OpenMEEG (Gramfort et al., 2010).

**Figure 2:**
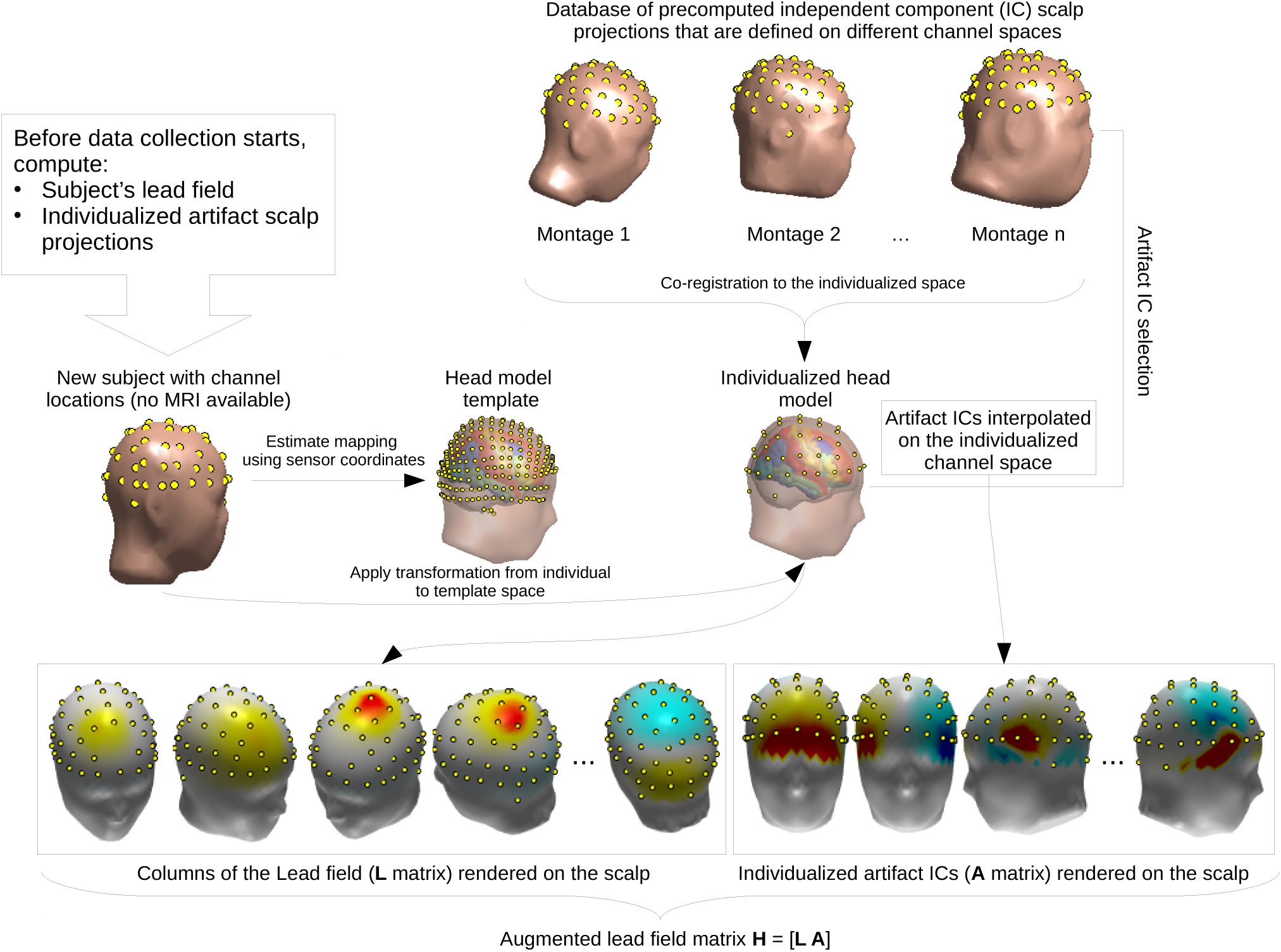
Procedure to obtain the lead field L and the dictionary of artifact scalp projections A.

Next, we identify the subjects in the database containing stereotypical artifact ICs (see next section) and co-register each of them with the individualized head model. We linearly map the artifact IC scalp projections to the sensor space of the participant using the transformation obtained during co-registration so that each warped IC has the same column-length as **L**, and can be appended to it. Finally, we divide each column of **H** by its norm so that the relative contribution to each source is determined by the amplitude of the source activation vector **x**_*k*_ only.

#### 2.1.2. Identification of artifact scalp projections

Several algorithms can be used to automatically identify stereotypical artifact scalp projections derived from ICA (Pion-Tonachini et al., 2017; Winkler et al., 2011). As a proof of principle, however, here we rely on the expertise of the authors. Since inspecting each IC in a large database is a cumbersome task, we propose the following simplification. We first co-register each subject in the database with the head model template to map ICs defined on different sensor spaces to a common space. We collect the mapped ICs in a matrix of 339 (number of channels of the template) by the number of total ICs. We reduce dimensionality by applying the k-means algorithm to the columns of the IC matrix. After inspecting the cluster centroids we label them as Brain, EOG, EMG, or Unknown (cluster of scalp maps of unknown origin). Finally, we store the indices of the EOG and EMG cluster centroid nearest neighbors for further use in the automatic creation of the **A** dictionary.

### 2.2. Spatiotemporal constraints

Since Eq (3) does not have a unique solution, to obtain approximated source maps with biological interpretation, we introduce constraints. One way of incorporating constraints in a principled manner is to express them in the form of the prior pdf of the sources *p*(**x**_*k*_). Since the neural generators of the EEG are assumed to be the electrical currents produced by distributed neural masses that become locally synchronized in space and time (Nunez and Srinivasan, 2006), here we seek to parameterized the prior *p*(**x**_*k*_) such that it induces source maps that are globally sparse (seeking to explain the observed scalp topography by a few spots of cortical activity) and locally correlated (so that we obtain spatially smooth maps as opposed to maps formed by scattered isolated sources) in space and time. Artifactual sources, on the other hand, can be assumed to be spatially uncorrelated from one another and from true brain sources.

A natural way to introduce the spatiotemporal constraints mentioned above into ESI is to model the source dynamics in the state-space framework. In this framework, Eq. (3) represents the observation equation and we assume the following state (source) transition equation,

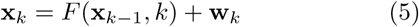

where the vector function *F* = [*F*_*g*_(·)^*T*^, *F*_*ν*_ (·)^*T*^]^*T*^ models how the source activity evolves from one sample to the next and **w**_*k*_ is a perturbation vector.

Several linear (Yang et al., 2016; Fukushima et al., 2015, 2012; Lamus et al., 2012; Cheung et al., 2010; Galka et al., 2004) and nonlinear (Olier et al., 2013; Giraldo et al., 2010; Valdes-Sosa et al., 2009; Daunizeau et al., 2009) models have been proposed for the brain state transition function *F*_*g*_(**g**_*k*_, *k*). For simplicity, here we use the linear model proposed by Yamashita et al. (2004) in which *F*_*g*_(**g**_*k*_, *k*) is reduced to a time-invariant linear operator describing the dynamics of the cortical activity due to nearby source interactions. In this approach, the evolution of the *i*th source *g*_*i,k*_ is given by the following first order autoregressive model,

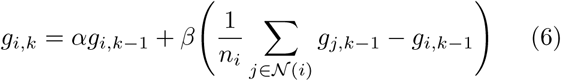

where the constants *α* and *β* are set to yield an observable system (Galka et al., 2004), 𝒩(*i*) contains the indices of the direct neighbors of dipole *i* and *n*_*i*_ is its total number of neighbors. Dipole neighbours can be extracted from the tessellation of the cortical surface. In the absence any other obvious transition model for artifact sources, we propose a simple random walk, which together with Eq. (6) yields the following linear transition function,

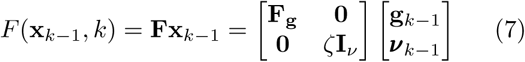

where the constant damping parameter *ζ* = 0.99 has the function of stabilizing the random walk model.

The state perturbation process **w**_*k*_ encompasses modeling errors as well as random inputs coming from distant sources, which we assume to be Gaussian and serially uncorrelated, **w**_*k*_ ∼ *N*(**w**_*k*_|**0, Q**_**k**_). We model the state noise covariance matrix **Q**_**k**_ with the following block-diagonal structure

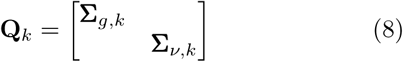

where the component of the covariance affecting brain sources are defined as follows,

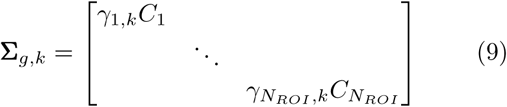

and 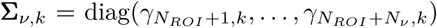 is the covariance of the noise component affecting artifact sources. We use this parameterization because it has been shown to induce group-sparse source estimates (Zhang and Rao, 2013). The matrices 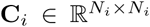 encode the intra-group brain source covariances and are precomputed based on source distance taking into account the local folding of the cortex as described in (Ojeda et al., 2018). 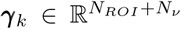 denotes a nonnegative scale vector that encodes the sparsity profile of the group of sources. Here we define *N*_*ROI*_ = 68 groups based on anatomical regions of interest (ROI) obtained from the Desikan-Killiany cortical atlas (Desikan et al., 2006). These assumptions together with the state transition model yield the following conditional source prior,

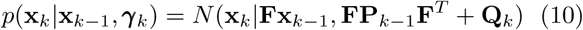

where **P**_*k*−1_ is the state covariance at *k* − 1 and we assume *p*(**x**_0_|**x**_−1_, *γ*_0_) = *N* (**x**_0_|**0, Q**_0_).

We complete our PGM by specifying priors on the hyperparameters *λ*_*k*_ and *γ*_*k*_ (hyperpriors). Assuming that *λ*_*k*_ and *γ*_*i,k*_ are independent yields the factorization

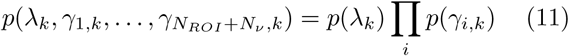

And since they are scale hyperparameters, we follow the popular choice of assuming Gamma hyperpriors with noninformative scale and shape parameters on a logscale of *λ*^−1^ and 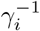 (Tipping, 2001). This choice of hyperprior has the effect of assigning a high probability to low values of *γ*_*i*_, which, in the static case (no transition equation), has been shown to shrink the irrelevant components of **x**_*k*_ to zero (Wipf and Nagarajan, 2008), thereby leading to a sparsifying behavior know as Automatic Relevance Determination (ARD) (Neal, 1996; MacKay, 1992).

We remark that although we are not the first to propose modeling the inverse solution of the EEG in the state-space framework, to our knowledge, we are the first to include artifact sources in the model and group-sparsity constraints. Fig. 3 shows the graphical representation of the proposed PGM.

**Figure 3:**
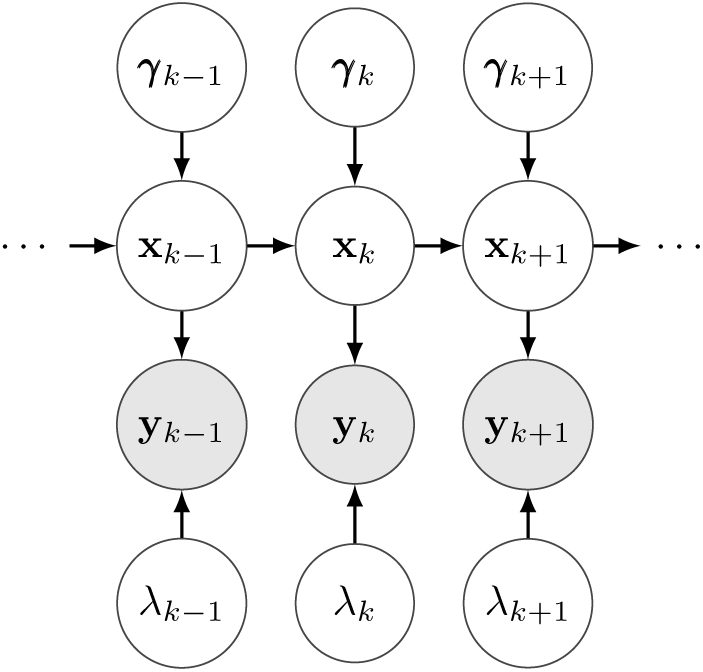
Graphical representation of the proposed generative model. Square, circle, and shaded circle symbols represent constant, hidden, and measured quantities respectively.

### 2.3. The Kalman filter

Our generative model belongs to the family of linear Gaussian dynamic systems (LGDS). In this type of systems, the source time series can be estimated from data optimally using the Kalman filter (Kalman, 1960). Because our algorithm is closely related to the Kalman filter, next we briefly outline this approach. To keep the notation uncluttered, in this section we remove the sample index *k* from the hyperparameters *λ* and *γ*.

Using the Bayes theorem we write the posterior of the sources as follows,

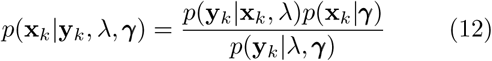

from which we have removed the dependency on the previous state, **x**_*k*−1_, by computing the predictive density

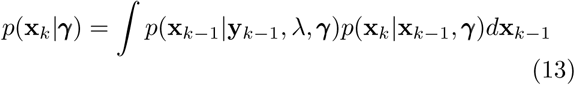

where *p*(**x**_*k*−1_|**y**_*k*−1_, *λ, γ*) is the source posterior already determined in the previous time step, *k* − 1. In a LGDS, the predictive density is also Gaussian with the following mean and covariance (Ozaki, 2012),

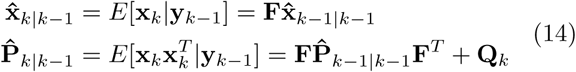

where we use the subindex *n*|*m* (with *n* ≥ *m*) to denote quantities estimated in the step *n* using data up to the *m* sample. Eq. (14) is known as the *time update*.

Since the numerator of Eq. (12) is the product of two Gaussian distributions, the posterior is also a Gaussian with the following mean and covariance (Ozaki, 2012),

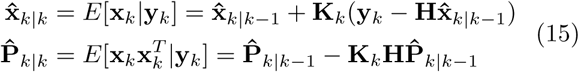

where **K**_*k*_ represents the Kalman gain,

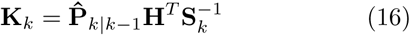

and **S**_*k*_ is the covariance of the data sequence **y**_*k*_. Eq. (15) is known as the *measurement update*. Using Eq. (14)-(16), we write the Kalman filter as the following recursive formula,

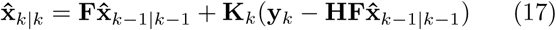

### 2.4. The RSBL filter

Yamashita et al. (2004) pointed out that the use of the Kalman filter in the context of ESI can be prohibitively expensive due to the necessity to estimate the *N*_*x*_ × *N*_*x*_ state covariance matrix **P**_*k*_. In that paper, the authors proposed that a tractable approximation to sidestep this problem is to remove the contribution of the neurodynamic model from the evolution of the state covariance, 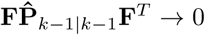, in Eq. (14). With this modification, we propose the use of the main recursive formula of the Kalman filter in Eq. (17) subject to the following gain and data covariance matrices,

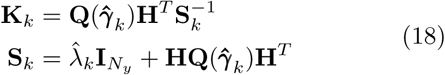

where 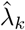 and 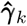 are optimal hyperparameter estimates obtained through the SBL algorithm outlined in Section 2.6. We summarize the RSBL filter in Algorithm 1 below.

#### Algorithm 1 RSBL filter

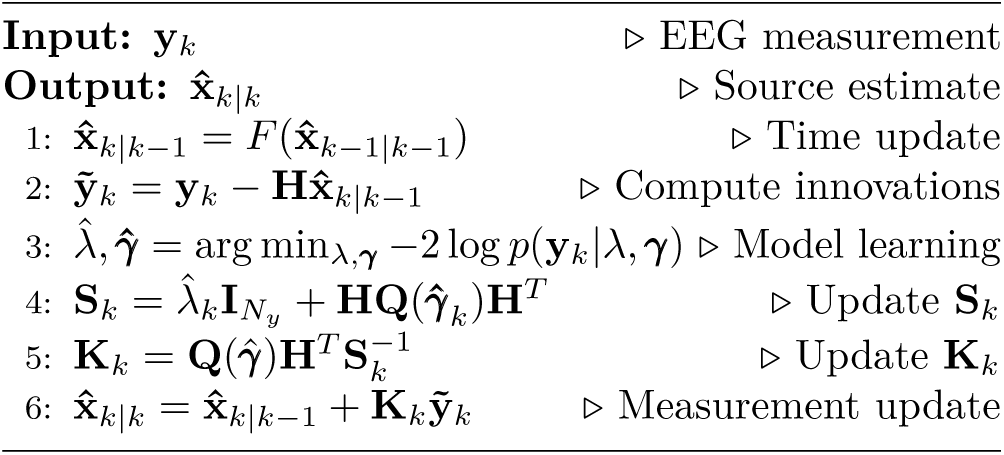

It is important to note that unlike most common use cases of the Kalman filter, in ESI we do not have access to a perfect description of the brain dynamics that is computationally tractable. Thus, we usually rely on approximate neurodynamic models, of which Eq. (6) is an example. We point out that our transition function is designed to provide a simple local spatiotemporal filtering effect that can be justified on the basis of nearby source interactions. However, the actual neurodynamic model may be far more complex. Since modeling errors are unavoidable, we argue that it makes sense from a practical standpoint to downplay the role of the neurodynamic model in the update of the state covariance. To compensate for this additional approximation, we exploit the block structure of the state vector induced by the hyperparameter vector *γ* to constrain the correction of the measurement update to regions of the state-space with biological significance.

Our approach differs from the one proposed by Yamashita et al. (2004) in two ways. First, they regularize the same amount everywhere in the source space (equivalent to considering a single *γ* parameter for the whole cortex) while we do so in groups of nearby sources encouraging sparsity in the ROI domain. Second, they do not consider time-varying hyperparameters and we do (*λ*_*k*_, *γ*_*k*_). Fixed hyperparameters may limit the applicability of this approach to experiments where the sparsity profile of the sources remains constant (e.g., steady state dynamics).

### 2.5. Data cleaning and source separation

The RSBL filter proposed above can be used to obtain a cleaned version of the EEG signal 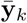, e.g. for visualization or scalp ERP analysis. To that end, we subtract the artifact components from the data as follows

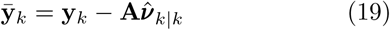

where 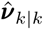 is obtained by selecting the last *N*_*ν*_ elements of the state vector 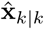.

Likewise, the estimated source vector ĝ_*k*|*k*_ is obtained by selecting the first *N*_*g*_ elements of 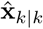. The source activity specific to the the *i*th ROI, 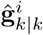, can be extracted from **ĝ**_*k*|*k*_ using the indices pulled from the cortical atlas. In some cases, further analysis of the source time series (e.g., source ERP and connectivity analysis) may be be carried out in the ROI space calculating the mean (or RMS mean if the dipole directions can be ignored) source activity within ROIs,

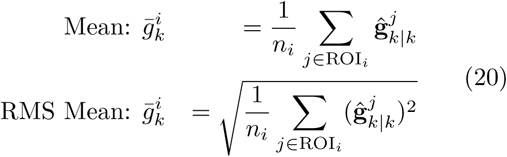

where ROI_*i*_ ⊂ {1, …, *N*_*g*_} is the subset of indices that belong to the *i*th ROI and *n*_*i*_ is the total number of sources within it.

### 2.6. Sparse model learning

The source estimates and cleaned data can be obtained analytically by evaluating the formulas given in Eq (17)-(20). These formulas, however, depend on the specific values taken by the hyperparameters that control our PGM, *λ*_*k*_ and *γ*_*k*_. In this section we outline the SBL algorithm for learning those.

The density function in the denominator of Eq. (12), *p*(**y**_*k*_|*λ*_*k*_, *γ*_*k*_), is known as the model evidence (MacKay, 2008b) and its optimization allows us to reshape our modeling assumptions in a data-driven manner. Since we used noninformative hyperpriors (see Section 2.2), to obtain source estimates conditioned on the optimal model we optimize the evidence,

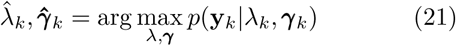

The evidence of a linear Gaussian model like ours is readily expressed as follows,

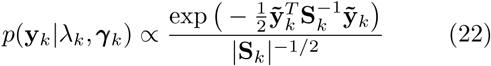

where 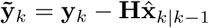 is the innovation sequence.

The maximization of the model evidence is equivalent to minimizing the Type-II Maximum Likelihood (ML-II) cost function, which is obtained by applying −2 log(·) to Eq (22),

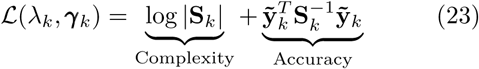

Eq (23) embodies a tradeoff between model complexity and accuracy. Geometrically, the complexity term represents the volume of an ellipsoid defined by **S**_*k*_. When the axes of the ellipsoid shrink due to the pruning of irrelevant sources, the volume is reduced. The second term is a squared Mahalanobis distance that measures model accuracy.

Eq (23) can be minimized very efficiently with a two-stage SBL algorithm proposed by Ojeda et al. (2018). In the first stage we learn a coarse-grained non-sparse model by solving the constrained optimization problem:

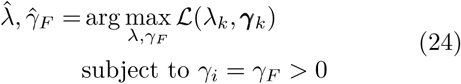

In the second stage we fix *λ* to the value 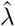 and, starting from the value 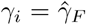, we learn the sparse model by solving the optimization problem:

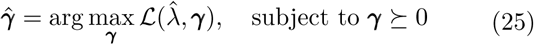

Intuitively, this process can be seen as an initial fast and reasonable, albeit coarse-grained, estimation, followed by a fine tuning step. We point the reader interested in the details of this approach to (Ojeda et al., 2018) while MATLAB code and examples can be freely downloaded from the *Distributed Source Imaging (DSI)* toolbox repository^1^.

### 2.7. Independent Component Analysis

In the analysis presented above, the matrix **H** is prespecified. In this section, we analyze the generative model of Eq. (3) from the ICA viewpoint. ICA is a blind source separation method that seeks to estimate the source time series (called activations in the ICA literature) **x**_*k*_ from the data time series **y**_*k*_ *without* knowing the gain (mixing) matrix **H**. In ICA, we assume that the latent sources are instantaneously independent, which yields the following prior distribution,

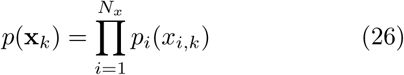

To simplify the exposition, we assume the same number of sensors and sources, *N*_*y*_ = *N*_*x*_, and the interested reader can find the case *N*_*y*_ < *N*_*x*_ in (Le et al., 2011; Lewicki and Sejnowski, 1998). From these premises, the objective of the algorithm is to learn the unmixing matrix **H**^−1^ such that we can estimate the sources with 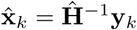. The unmixing matrix **Ĥ** ^−1^ can be learned up to a permutation and rescaling factor, which has the inconvenience that the order of the learned components can change depending on the starting point of the algorithm and data quality. We can use a data block **Y** to write the likelihood function,

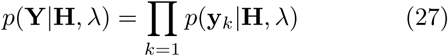

under the assumption of independent data collection. We should point out that in the case of EEG, the signal is not iid because of the short term autocorrelations produced by the underlying source dynamics. To alleviate this situation the data are usually whitened during preprocessing. We obtain each factor in Eq (27) by integrating out the sources as follows,

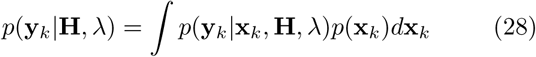

As noted by MacKay (2008a), assuming that the data are collected in the noiseless limit, *λ* → 0, transforms the Gaussian likelihood *p*(**y**_*k*_ |**x**_*k*_, **H**, *λ*) into a Dirac delta function, in which case Eq (28) leads to the Infomax algorithm of Bell and Sejnowski (1995). The learning algorithm essentially consists in finding the gradient of the log likelihood, log *p*(**Y**|**H**, *λ*), with respect to **H** and updating **H** on every iteration such that the likelihood of the data increases. As pointed out by Comon (1994), the ICA model is uniquely identifiable only if at most one component of **x**_**k**_ is Gaussian. Therefore, the prior densities *p*_*i*_(*x*_*i,k*_) are usually assumed to exhibit heavier tails than the Gaussian and, in particular, the prior *p*_*i*_(*x*_*i,k*_) ∝ cosh^−1^ *x*_*i,k*_ yields the popular ICA contrast function tanh(**H**^−1^**y**_*k*_). Note that this prior is not motivated by a biological consideration but by a mathematical necessity.

It is remarkable that ICA can learn columns of **Ĥ** that are consistent with bipolar (single or bilaterally symmetric) cortical current source scalp projections without using any anatomical or biophysical constraint whatsoever (Makeig et al., 1997). Onton et al. (2006) have shown that other columns may correspond to different stereotypical artifact scalp projections as well as a set of residual scalp maps that are difficult to explain from a biological standpoint. Delorme et al. (2012) have shown that the best ICA algorithms can identify approximately 30% of dipolar brain components (approximately 21 brain components out of 71 possible in a 71-channel montage). Although ICA has proven to be a useful technique for the study of brain dynamics (Makeig and Onton, 2011), we must wonder if its performance can be improved, perhaps by making BSS of EEG data less “blind”. In other words, if we know a priori what kind of source activity we are looking for (dipolar cortical activity, EOG and EMG artifacts and so on), why limit ourselves to a purely blind decomposition?

In this paper, we advocate the use of as much information as we can to help solve the ill-posed inverse problem. In that sense, the use of a prespecified lead field matrix in the generative model of the EEG forces inverse algorithms to explain the data in terms of dipolar sources, because the lead field is precisely an overcomplete dictionary of dipolar projections of every possible source there is in a discretized model of the cortex. It has been shown that source estimation can greatly benefit from the use of geometrically realistic subject-specific (Cuspineda et al., 2009) or, alternatively, population-based approximated lead fields matrices (Valdés-Hernández et al., 2009). Furthermore, augmenting the lead field dictionary with a set of stereotypical artifact projections, as proposed in Section 2.1, furnishes a more realistic generative model of the EEG in a way that renders blind decomposition unnecessary or at least suboptimal for brain imaging.

### 2.8. Independent source separation through RSBL

It has been pointed out that, due to the volume conduction effect, the source estimates obtained from EEG are still a mixture of the actual source activity (Biscay et al., 2018). To guard against this problem, ideally, we would like the RSBL algorithm to exhibit the ICA property of yielding maximally independent (demixed) source time series. Recently, Biscay et al. (2018) used arguments complementary to those given in this section to show that, in the static case, the use of source ROI constraints induces the desired unmixing effect. Next, we show that this is indeed the case when source dynamics are considered and that it is a consequence of parameterizing the sources with sparse priors. We start by rewriting the biologically motivated source prior of Eq (13) as follows,

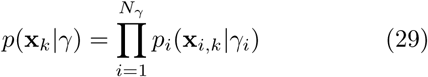

where each factor is a Gaussian pdf and *i* indexes a group of sources or an artifact component. To write Eq (29) as the ICA prior in Eq (26) we need to integrate out the hyperparameter *γ*_*i*_ from each factor as follows:

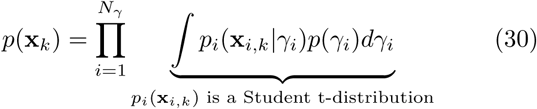

which, given our choice of hyperprior on *γ*_*i*_, renders each marginalized prior *p*_*i*_(**x**_*i,k*_) a heavy-tailed Student t-distribution (Tipping, 2001). We note that in our development we take the route of optimizing the *γ*_*i*_ hyperparameters rather than integrating them out because the former approach yields a simpler algorithm and tends to produce more accurate results in ill-posed inverse problems (MacKay, 1996). The optimization of *γ*_*i*_ allows for automatic removal of irrelevant brain and artifact components that are not supported by the data, thereby eliminating the subjectivity implicit in manual component selection. Assuming the prior in Eq (29), the ICA data likelihood of Eq (28) becomes exactly the evidence of Eq (22), with the difference that in the RSBL filter the **H** matrix is known and the evidence is optimized on every sample, which gives our algorithm the ability to run online and to capture transient dynamics.

We summarize the advantages of using the RSBL filter over ICA for source separation and imaging of EEG data as follows:

- First and foremost, artifact removal, source separation, and imaging can be obtained simultaneously as a consequence of optimizing the evidence of a biologically informed generative model.
- It deals gracefully with the overcomplete case (*N*_*y*_ ≪ *N*_*x*_) by finding a regularized source estimator, which always exists even in the presence of rank-deficient data, e.g., after removing the common average reference.
- It deals with the redundancy in brain responses by inducing independence over groups of sources.
- The use of the ARD prior allows for the automatic selection of components in a data-driven manner.
- It can adapt to non-stationary dynamics by updating the model on every sample.
- It can be used in online applications by leveraging fast evidence optimization algorithms.
- It facilitates subject-level analysis because we estimate the same number of cortical source activations per subject, each of which has known anatomical support. This eliminates the complications of clustering ICs and dealing with missing components (Bigdely-Shamlo et al., 2013b) while allowing the use of more straightforward and widespread statistical parametric mapping techniques (Penny et al., 2007).

## 3. Results

In this section we first describe the data used to obtain the artifact dictionary. In Section 3.2 we test the RSBL algorithm on simulated data. In sections 3.3-3.7 we test the algorithm on real data.

### 3.1. Empirical characterization of artifact scalp projections

To construct the artifact dictionary we used data from two different studies made public under the umbrella of the BNCI Horizon 2020 project^2^ (Brunner et al., 2015). We briefly describe these data sets next.

#### Data set 1: Error related potentials

The first study, *013-2015*, provided EEG data from 6 subjects (2 independent sessions per subject and 10 blocks per session) collected by Chavarriaga and del R. Millán (2010) using an experimental protocol designed to study error potentials (ErrP) during a BCI task. EEG samples were acquired at a rate of 512 Hz using a Biosemi ActiveTwo system and a 64-channels montage placed according to the extended 10/20 system.

#### Data set 2: Covert shifts of attention

The second study, *005-2015*, provided EEG and EOG data from 8 subjects collected by Treder et al. (2011) using an experimental protocol designed to study the EEG correlates of shifts in attention. The EEG was recorded using a Brain Products actiCAP system, digitized at a sampling rate of 1000 Hz. The montage employed had 64 channels placed according to the 10/10 system referenced to the nose. In addition, an EOG channel (labeled as EOGvu) was placed below the right eye. To measure vertical and horizontal eye movements, from the total of 64 EEG channels, two were converted into bipolar EOG channels by referencing Fp2 against EOGvu, and F10 against F9, thereby yielding a final montage of 62 EEG channels.

#### Data preprocessing and IC scalp map clustering

After transforming each data file to the .*set* format, both studies were processed using the same pipeline written in MATLAB (R2017b The MathWorks, Inc., USA) using the EEGLAB toolbox (Delorme et al., 2011). The pipeline consisted of a 0.5 Hz high-pass forward-backward FIR filter and re-referencing to the common average, followed by the Infomax ICA decomposition of the continuous data. We pooled all the preprocessed files from all subjects in both data sets and randomly assigned them to one of two groups: 80 % to the training set and 20 % to the test set. The training set was used to construct the EEG database and the artifact dictionary while the test set was used to evaluate the performance of the RSBL algorithm on real data in subsections 3.3-3.5. Note that in this approach, we use the testing set to simulate new subjects whose artifacts are not explicitly characterized in the database. Finally, we calculated the lead field of the subjects in the testing set as described in Section 2.1.1.

To select artifact ICs we used the training set and followed the procedure outlined in Section 2.1.2. After performing the co-registration to the template, the training set resulted in a matrix of 339 channels by 6774 independent scalp maps (101 sessions and blocks yielding 64 ICs each plus 5 sessions yielding 62 ICs each). Fig 4 shows a visualization of the IC scalp maps using the t-distributed stochastic neighbor embedding (t-sne) algorithm (Van Der Maaten and Hinton, 2008). The t-sne algorithm allows us to represent each 339-dimensional IC scalp map as a dot in a 2D space in a way that similar and dissimilar scalp maps are modeled by nearby and distant points respectively with high probability. We ran the k-means algorithm with several numbers of clusters stopping at 13 after noticing that many small islands scattered at the periphery of Fig 4 started to be either mislabeled as Brain or labeled consistently as EOG, EMG or Unknown. Unknown clusters were not further used in this paper. The grey points in the figure denote most of the scalp maps labeled as non-brain.

**Figure 4:**
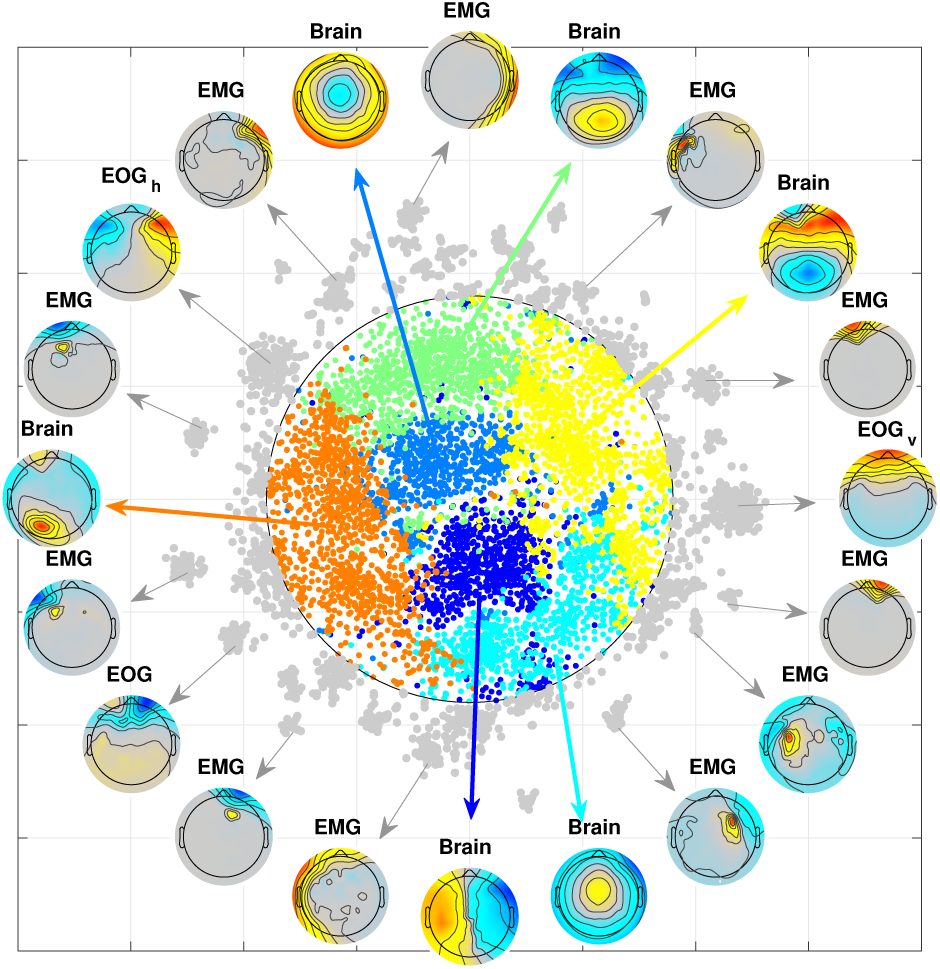
t-sne visualization of IC scalp map clusters. We used the t-sne algorithm to represent each 64-dimensional scalp map as a dot in a 2D space in a way that similar and dissimilar scalp maps are modeled by nearby and distant points respectively with high probability. The clusters were estimated using the k-means algorithm. The grey points indicate mostly non-brain or mislabeled scalp projections.

After the artifact selection and co-registration processes, we obtained the following **A** dictionary for each individualized head model in the testing set:

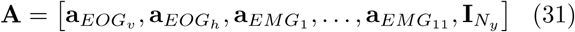

where 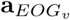 and 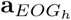 are the respective scalp projections of the vertical and horizontal EOG ICs, 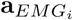 are the projections of 11 representative EMG ICs, and we modeled spike artifacts affecting each individual channel with the columns of the identity matrix 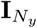.

### 3.2. Performance on simulated data

We used the time series of sources in two ROIs to simulate 2 seconds of evoked EEG data contaminated by the activity of two artifact sources (see Fig. 5). The objective of this experiment was to unmix the EEG signal and recover all four sources under different common-mode noise conditions. To that end, we placed brain sources in the anterior (ACC) and posterior (PCC) cingulate cortex. We designed their time courses to simulate the distribution of scalp voltages observed during ErrP studies (Gehring et al., 2012). ErrP are characterized by a negativity (Ne component) in the frontocentral channels within 100 msec after an erroneous response followed by a positivity (Pe component) in the centroparietal channels around the 250 msec. The timing of these components can be reliably identified by inspecting the EEG activity of frontocentral channels (Fz, FCz, Cz) time-locked to the erroneous responses.

**Figure 5:**
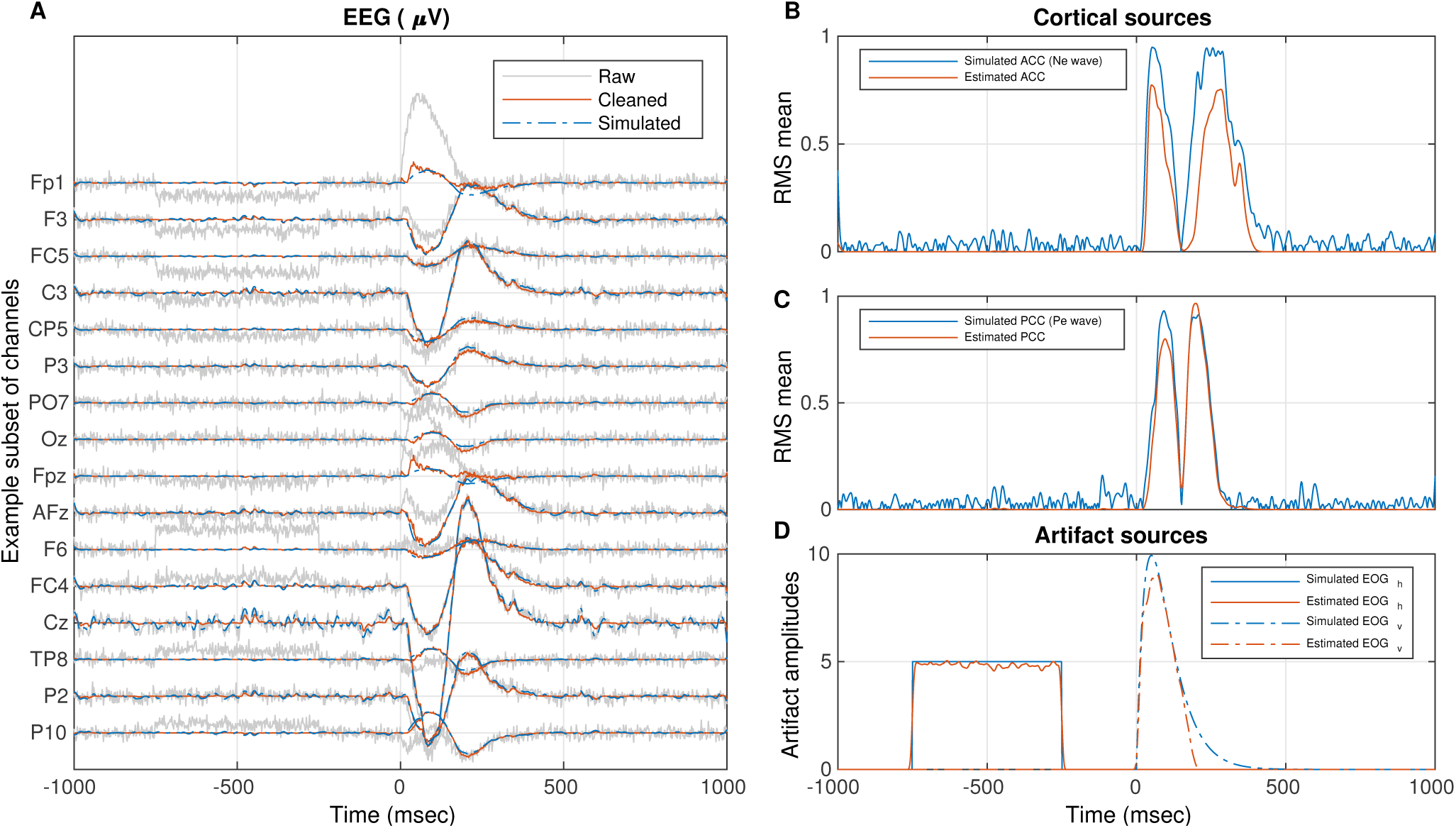
Example of RSBL spatiotemporal filtering on simulated data for a common mode SNR = 6 dB. **A**: Raw, simulated and estimated scalp time series for a subset of channels. **B, C, D**: Simulated and estimated time series of the magnitude of different cortical and artifact sources. The common mode SNR was defined with respect to the projection of the true cortical source activity before EOG artifacts were introduced. A common mode SNR of 6 dB indicates that that the amplitude of the simulated EEG signal was, on average, twice higher than the amplitude of the sensor noise.

We simulated Ne and Pe spatial profiles with the column vectors **g**_A_ and **g**_P_, which contained 1 only in the entries corresponding to dipoles in ACC or PCC respectively and 0 elsewhere. The temporal profiles, **t**_*Ne*_ and **t**_*Pe*_ were simulated using Gamma functions. We simulated the cortical activity by multiplying the spatial and temporal profiles as follows,

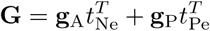

The two artifact sources simulated a lateral eye movement (−750 msec to 250 msec) and eye blink (0 msec to 400 msec) events. All other artifact sources were set to 0. The ratios between the maximum amplitude of artifacts, EOG_h_ and EOG_v_, to cortical source activity were set to 5 and 10 respectively. The simulated source activity 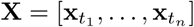 was generated by concatenating the cortical **G** and artifact source matrices. Scalp data was generated by projecting **X** to the sensors and adding white Gaussian noise. The variance of the noise was calculated with respect to the variance of the sensor data before adding EOG artifacts. We simulated the sensor space with a 64-channels montage placed according to the extended 10/20 system and computed **H** as described in Section 2.1.1.

Table 1 summarizes the performance of the RSBL algorithm for different levels of noise. Column one shows the SNR in dB units and column two shows the equivalent EEG signal to noise amplitude ratio (AR). The columns headed by ACC, PCC, EOG_v_, and EOG_h_ report the correlation value between their respective simulated and estimated sources; high values indicate good estimation accuracy. The columns headed by \ACC and \PCC report the mean correlation between simulated ACC and PCC sources and every other estimated cortical source; low correlation values indicate low source leakage. We note that for extremely aggressive noise conditions (i.e., SNR values between −10 dB and 0 dB) the algorithm failed to consistently reconstruct all the simulated sources accurately. However, once the amplitude of the EEG was at least one and a half or higher than the amplitude of the noise, we obtained correlations in the 0.63-0.99 range. In all cases, the leakage was low as indicated by correlation values below 0.04.

**Table 1:** RSBL performance for different SNR values. The columns ACC and PCC report the correlation between the simulated and the estimated sources of each respective ROI. The grayedout columns with the \ symbol in the header report the mean correlation between the simulated sources and all other estimated sources removing the respective ACC or PCC ROI; low values in this columns indicate low source leakage. The last two columns report the correlation between simulated and recovered artifact sources.

| SNR<sup>1</sup> (dB) | AR<sup>2</sup> | ACC | \ACC | PCC | \PCC | EOG<sub>v</sub> | EOG<sub>h</sub> |
| --- | --- | --- | --- | --- | --- | --- | --- |
| -10 | 0.316 | 0.0032 | 0.0200 | 0.016 | 0.0241 | 0.096 | 0.099 |
| -6 | 0.5 | 0.0200 | 0.0382 | 0.765 | 0.0327 | 0.109 | 0.111 |
| 0 | 1 | 0.1678 | 0.0189 | 0.964 | 0.0078 | 0.715 | 0.496 |
| 3 | 1.4 | 0.6325 | 0.0175 | 0.921 | 0.0175 | 0.739 | 0.803 |
| 6 | 2 | 0.8808 | 0.0129 | 0.869 | 0.0137 | 0.881 | 0.949 |
| 10 | 3.16 | 0.7211 | 0.0131 | 0.991 | 0.0098 | 0.969 | 0.983 |
| 20 | 10 | 0.8805 | 0.0121 | 0.993 | 0.0109 | 0.974 | 0.998 |

1. Signal to noise ratio. 2. EEG signal to amplitude ratio.

Fig. 5 illustrates the simulated data and results for SNR=6 dB. The traces in panel **A** represent the raw, cleaned, and simulated EEG time series for a subset of channels. The cleaned EEG traces were obtained by subtracting out the estimated EOG activity. The panels on the right show the ground truth and estimated source activity in the ACC (**B**), PCC (**C**) areas, as well as EOG artifacts (**D**). Note that the estimated source time series are sparse in space and time (see also Fig. 6). In particular, panels **B** and **C** demonstrate that source values that randomly oscillate at the noise level are ignored by the algorithm. That does not mean that those cortical areas are silent, but that the postsynaptic potentials produced by the collective firing of pyramidal cells in those places are not coherent enough to create signals that can be reliably measured by the scalp sensors.

**Figure 6:**
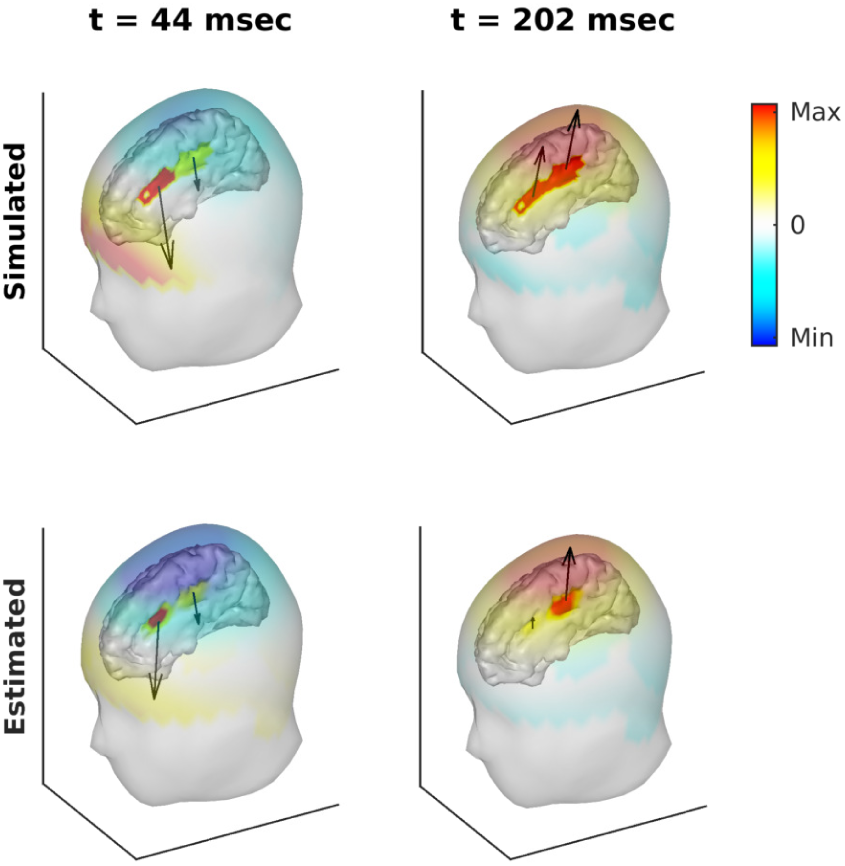
Simulated (top row) and estimated (bottom row) scalp and cortical maps at the peak of the Ne (44 msec) and Pe (202 msec) waves. We applied transparency to the head surfaces so that the cortical maps can be seen in the interior layer. The cortical maps show a view of the right hemisphere, exposing the activations in the cingulate gyrus. The color of the scalp maps represent EEG voltages. The color of the cortical maps represent the magnitude of the PCD at every cortical location. The arrows represent the average direction of dipoles inside each ROI. In the bottom row, we have removed the contribution of the estimated EOG activity to the scalp EEG.

Fig. 6 shows a snapshot of the scalp and source activities at the peak of the Ne and Pe components displayed in Fig. 5. We linearly extrapolated the sensor values to the portion of head covered by the EEG cap, so that the topographic maps could be rendered on the 3D surface. In the top row, we see that although all the simulated sources within an ROI have the same amplitude, in the recovered maps (bottom row) the nontrivial dipole PCD values are not identical (though correlated). The panels in the bottom row show the EEG scalp maps with the artifact activity subtracted out. Note that even when the scalp data is severely affected by the eye blink artifact at the peak of the Ne component (t = 44 msec), the algorithm correctly estimated the orientation and location of its generators in the ACC.

### 3.3. Single-trial analysis on real data

In this section, we demonstrate RSBL single-trial source analysis on real data. To that end, we selected a subject from the testing set that belonged to the ErrP study (data set 1 above). In the experimental protocol used by Chavarriaga and del R. Millán (2010), the subjects were tasked with moving a cursor towards a target location either using a keyboard or mental commands. The authors of that paper found consistent evoked potentials produced by errors induced by the computer. In other words, subjects elicited error potentials after watching the computer execute the wrong moves; this is sometimes referred to as feedback error-related negativity/positivity (Ward et al., 2013). These error potentials were characterized by two frontocentral positive peaks around 200 msec and 320 msec after the feedback, a frontocentral negativity around 250 msec, and a broader frontocentral negative deflection about 450 msec.

Fig. 7 summarizes the results of applying the RSBL filter to a single-trial of the experiment. Panel **A** shows the EEG signal of three frontocentral sensors (Fz, FCz, and Cz). Panel **B** shows the source magnitude time series averaged within the ACC and PCC regions, **C** shows the scalp and cortical maps at the latency of the Ne and Pe components, and **D** shows the maximum intensity projection of each source map in **C** in the sagittal plane. Panels **B** and **C** reveal that the error components observed at the scalp level are mostly generated by a complicated interplay of sources in the ACC and PCC regions. The sagittal maximum intensity projections, however, indicate that other cortical sources also contribute to the observed EEG signal, although a lesser amount. This is common in single-trial analyses, and usually, those sources that are not related to errors clear out after trial averaging. We note that the result that both ACC and PCC sources contribute to Ne and Pe scalp topographies is in agreement with recent experimental findings (Buzzell et al., 2017).

**Figure 7:**
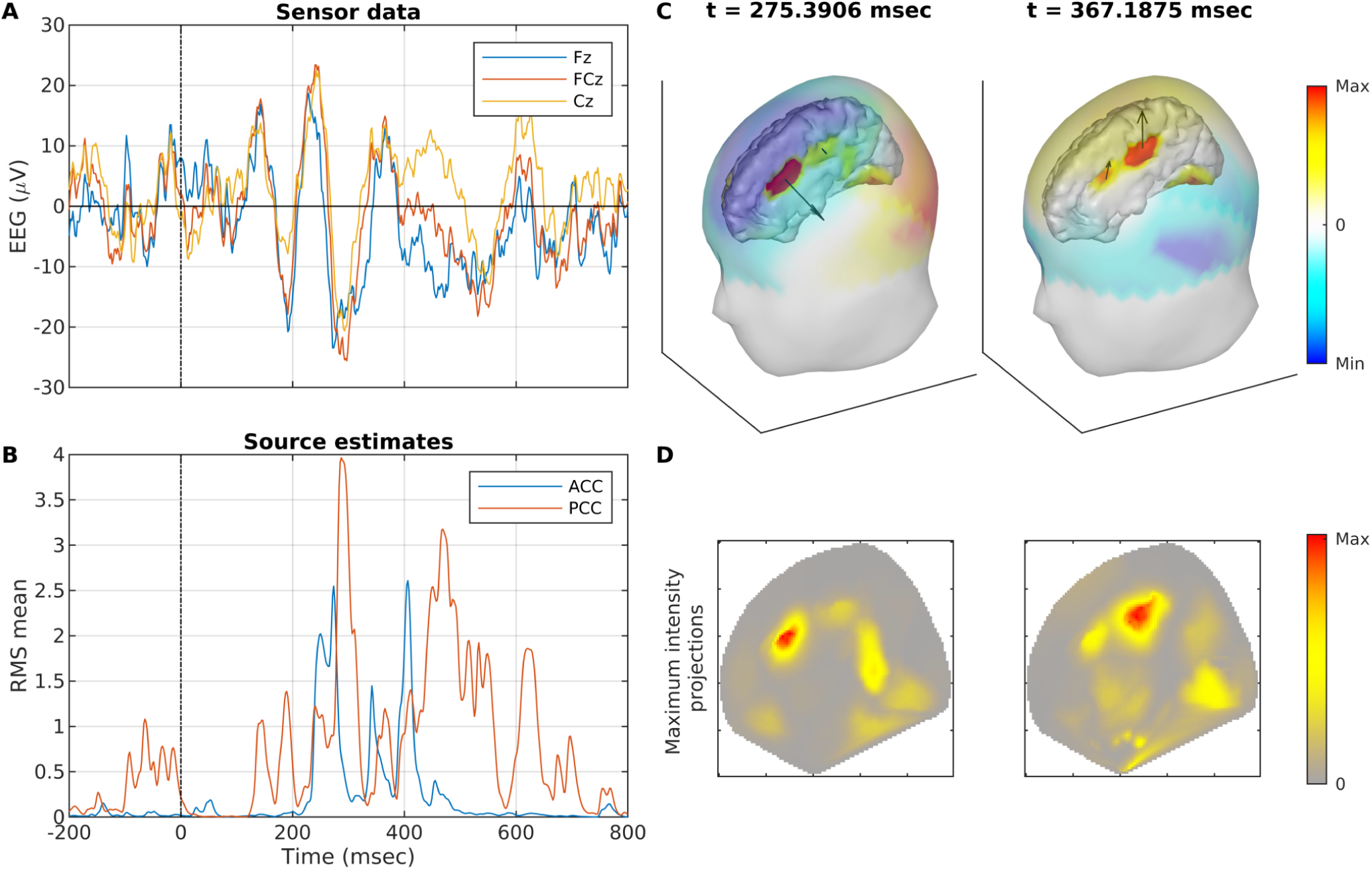
Example of RSBL single-trial source analysis. **A**: EEG data of frontocentral channels. **B**: Source magnitude times series averaged within ACC and PCC regions. **C**: EEG scalp maps and cortical estimates at the peak of the Ne and Pe components. **D**: Maximum intensity projection maps of the cortical activations in panel **C** along the sagittal plane.

### 3.4. EOG artifact removal on real data

In this example, we applied the RSBL algorithm to a trial contaminated by a lateral eye movement and eye blink activity. We used data from the same subject selected for the experiment in the previous section, but this time we analyzed an epoch with no error-related activity so that we could appreciate the artifact rejection performance of the algorithm minimizing task-related confounds. Fig. 8 summarizes the data and results.

**Figure 8:**
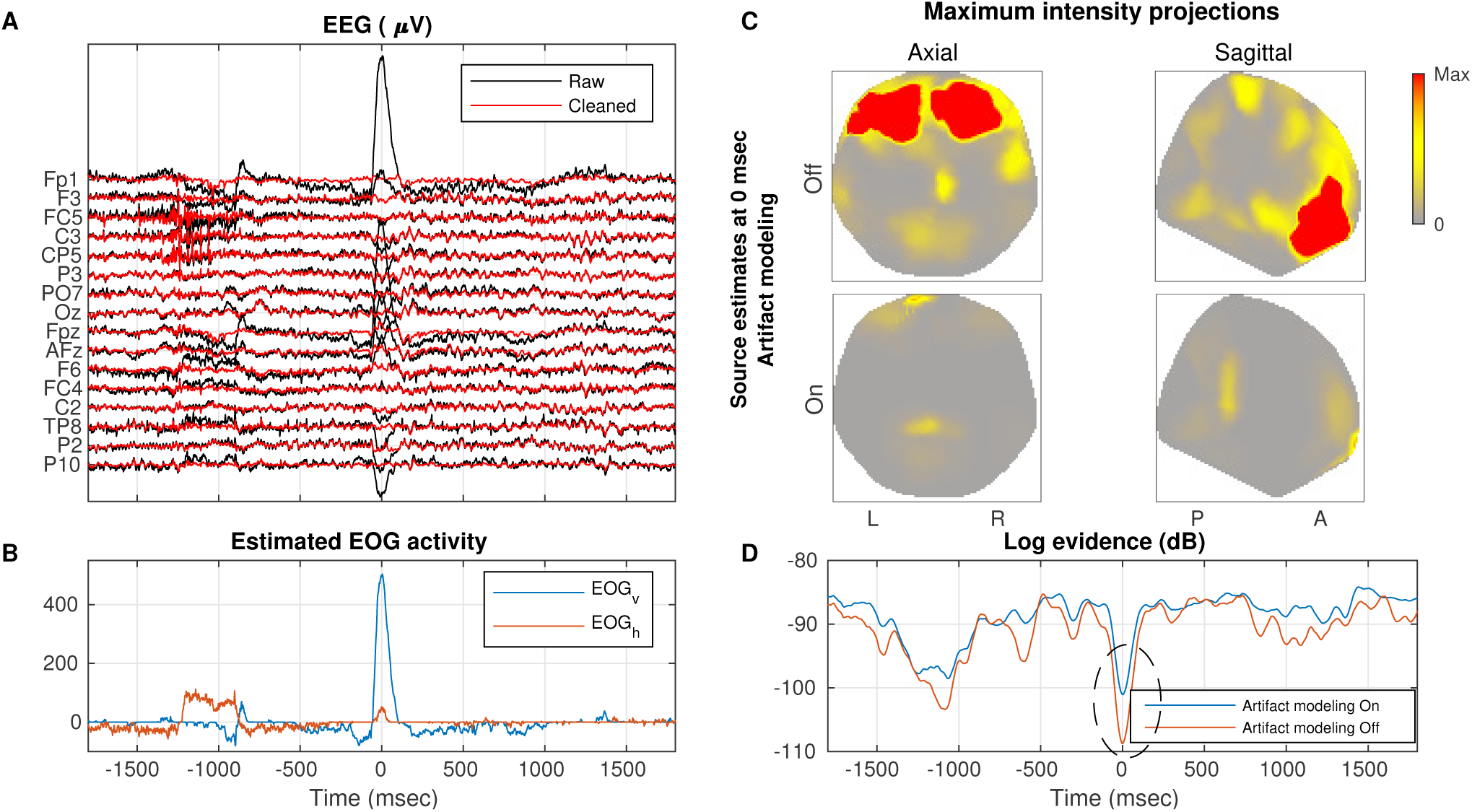
Example of RSBL artifact cleaning of an EEG epoch with lateral eye movement and eye blink artifacts. **A**: Subset of raw and cleaned EEG channels. **B**: Estimated EOG_*v*_ and EOG_*h*_ artifact source activity. **C**: Maximum intensity projection maps of the source map at the latency of the peak eye blink artifact with artifact modeling turned off (top row) and on (bottom row). **D**: Log evidence time series obtained by the algorithm with artifact modeling turned off and on.

In Fig. 8, panel **A** shows a subset of the raw and reconstructed (cleaned) EEG traces in black and red respectively. There is a lateral eye movement artifact between −1250 msec and −800 msec and an eye blink between −250 msec and 1000 msec. Panel **B** shows the estimated EOG_*v*_ and EOG_*h*_ artifact source activity in blue and orange, respectively. We note that these artifact sources were active at the latencies where the EEG is affected and mostly zero elsewhere. This is a desired feature of the algorithms because this way it only “fixes” the affected segments while leaving clean data unchanged (this result is generalized to several subjects and epochs in Section 3.5). In **C** we show the maximum intensity projection maps of the source map at the latency of the peak eye blink artifact, 0 msec. Each column displays a different projection. The rows display source estimates without and with artifact modeling enabled. Panel **D** shows the time series of the log evidence for generative models with artifact modeling on and off. In panel **D**, both traces differ mostly only when artifacts occur, where higher log evidence of the blue trace indicates that source estimation benefits from modeling artifacts. This is not a striking finding, but it illustrates the practical utility of the evidence metric for data modeling.

Visual inspection of the bottom row of panel **C** reveals that some residual eye blink activity may have leaked into the frontal pole. We point out that, in practice, it is extremely hard to totally remove artifactual activity because: 1) the use of a lead field matrix derived from a template head model may misfit the anatomy of the subject, introducing errors in the **L** dictionary, 2) errors in the sensor locations can cause the EEG topography to shift with respect to the expected brain and artifact source projections, 3) EMG scalp projections are difficult to characterize due to their variability, as opposed to EOG projections that are more stereotyped, and 4) unmodeled artifactual activity, such as muscle projections towards the back of the head that were largely ignored in this study, may be suboptimally accounted for. Despite all these issues, Fig. 8 demonstrate that RSBL can yield reasonably robust source estimates in the presence of high amplitude artifacts. Furthermore, panel **D** suggests that we could use dips in the log evidence to inform subsequent processing stages of artifactual events that were not successfully dealt with.

### 3.5. Data cleaning performance: benchmark against ASR

Next, we benchmarked the data cleaning performance of the RSBL algorithm against ASR. The ASR algorithm has gained popularity in recent years for its ability to remove a variety of high amplitude artifacts in an unsupervised manner, thereby enabling automatic artifact rejection for offline as well as real-time EEG-based BCI applications. Since in real data we do not have a ground truth for artifactual activity, we compare both methods according to the correlation between raw and cleaned data samples in blocks with negligible or no artifactual activity, where low correlation values indicate needless distortion of the brain activity.

We ran both algorithms for each subject in the test set and collected the following quantities on subsequent blocks of 40 msec: 1) the correlation between raw and cleaned data (computed as the correlation between the correspondent data blocks vectorized across channels and time points) and 2) the maximum RMS artifact power yielded by RSBL. ASR’s performance depends on multiple parameters, but it has been shown that the most critical one is the cutoff (Chang et al., 2018). In this experiment we used a cutoff equal to 5, which was the default value of EEGLAB’s ASR plugin at the time of preparing this publication.

In Fig 9, the left and right panels show the empirical kernel pdf estimation of the correlation as a function of the artifact’s power for the ASR and RSBL algorithms, respectively. We see that in both methods, the correlation decreases as artifact power increases. This effect is expected and desired because cleaning algorithms are supposed to modify contaminated raw data. Towards low power artifact regions, however, ASR exhibits a significant amount of probability mass that spreads down to low correlation values while RSBL seems to have most of its probability mass bounded from below at around 0.8. This result indicates that, at a cutoff of 5, ASR cleaning is overly aggressive to the point of significantly modifying the data in the absence of artifacts. These findings are in agreement with what was recently reported by Chang et al. (2018). In that paper, the authors determined that the optimal cutoff parameter of ASR may be between 10 and 100. We remark, however, that a practical advantage of RSBL over ASR is that in the former, all the parameters are automatically learned from the data thereby removing the need for user intervention or calibration.

**Figure 9:**
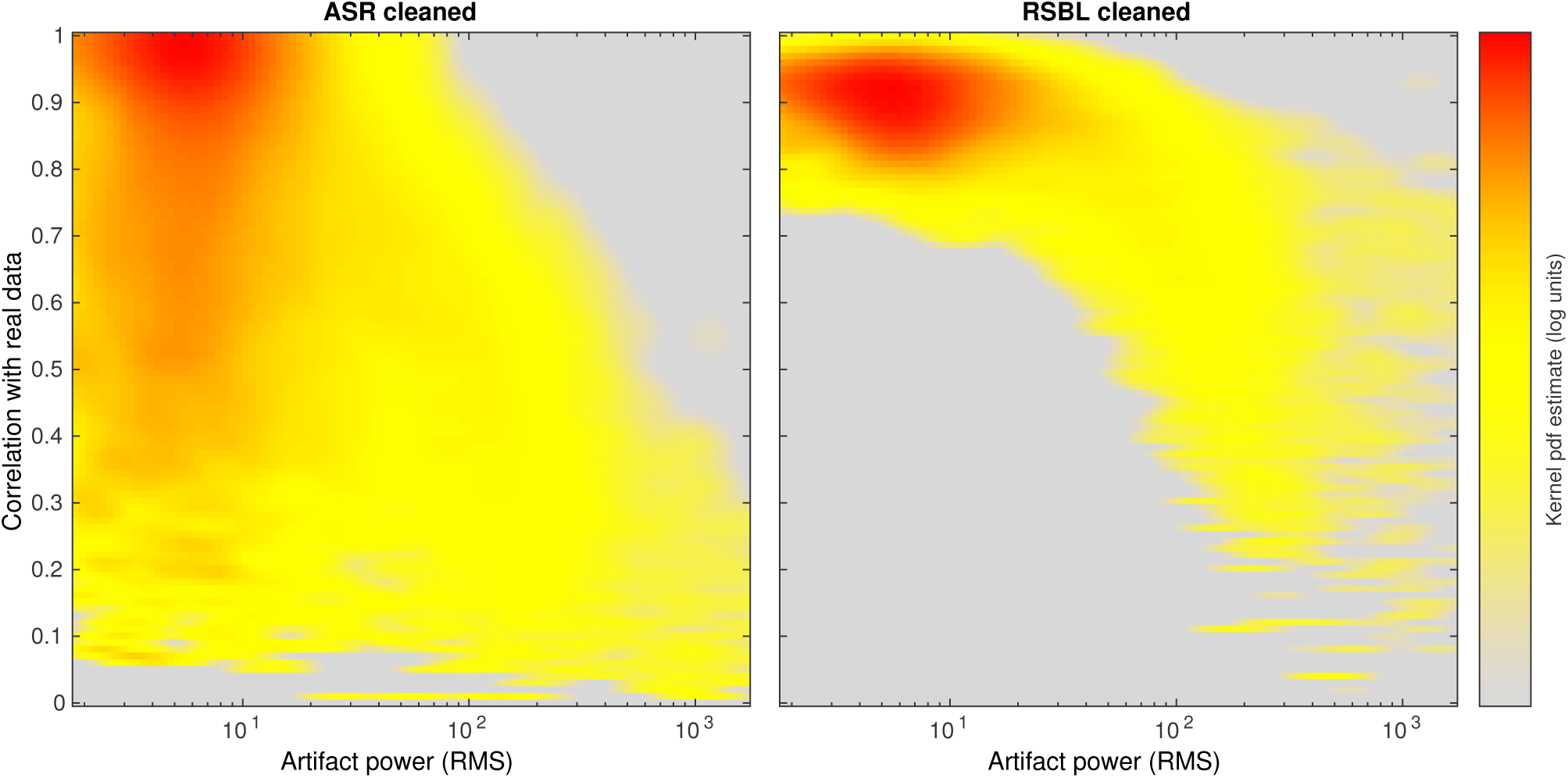
Data cleaning performance. Kernel pdf estimation of the correlation between raw and cleaned data as a function of artifact power. **Left:** Data cleaned by ASR using cutoff=5 (default). **Right:** Data cleaned by RSBL. Note that, as expected, in both algorithms the correlation drops as artifacts increase. Towards low amplitude artifacts, however, ASR significantly distorts the data while RSBL does not.

### 3.6. Source separation performance: benchmark against Infomax ICA

In this section, we investigated the source separation performance of the RSBL algorithm. For this, we used the test set to benchmark RSBL against Infomax ICA regarding volume conduction unmixing as a function of the data size. We assessed the unmixing performance by calculating the mutual information reduction (MIR) achieved by each algorithm on data blocks of increasing sizes.

The MIR is an information theoretic metric that measures the total reduction in information shared between the components of two sets of multivariate time series. The mutual information (MI) between two given time series *x*_*i,k*_ and *x*_*j,k*_, *I*(*x*_*i*_, *x*_*j*_), can be defined as the Kullback-Leibler (KL) divergence between their joint and marginal distributions:

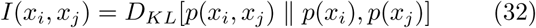

where *I*(*x*_*i*_, *x*_*j*_) > 0 indicates that processes *x*_*i*_ and *x*_*j*_ share information while *I*(*x*_*i*_, *x*_*j*_) = 0 indicates that they are statistically independent such that

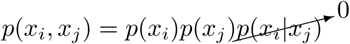

We define the MIR of source separation algorithm A with respect to B, as the difference in normalized total pairwise MI (PMI) achieved by each decomposition:

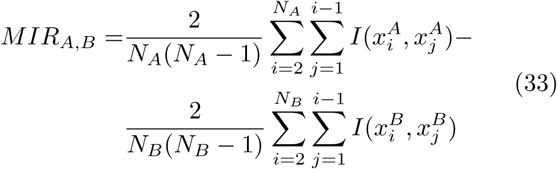

where 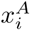 and 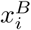 are the set of components yielded by each method and *N*_*A*_ and *N*_*B*_ are the number of components afforded by each decomposition. We note that to obtain a PMI that is not biased by the number of components, we normalize each summation by the number of unique (*i, j*) pairs. Here we calculated the MI using the empirical estimates of the distributions in Eq (32) for the the multichannel EEG data, the ROI-collapsed sources estimated by RSBL, and the ICs obtained by Infomax.

In Fig 10, the left panel shows a box plot of the MIR of RSBL and Infomax calculated with respect to the MI of channel data. As indicated by the x-axis, we ran the experiment multiple times increasing the data sizes from 2 to 500 seconds (∼ 8 minutes). As expected, both algorithms reduced source MI, thereby reversing to some extent the mixing effect of the volume conduction. We see also that, on average, when the MIR is calculated in short blocks of data, RSBL exhibited higher unmixing performance while Infomax did better on longer blocks. This effect is more clearly represented in the panel on the right, which shows the box plot of the MIR of RSBL with respect to Infomax. In that panel, distributions with entire positive (orange) or negative (blue) values indicate a significant source crosstalk reduction performance in favor of the RSBL or Infomax algorithms respectively.

**Figure 10:**
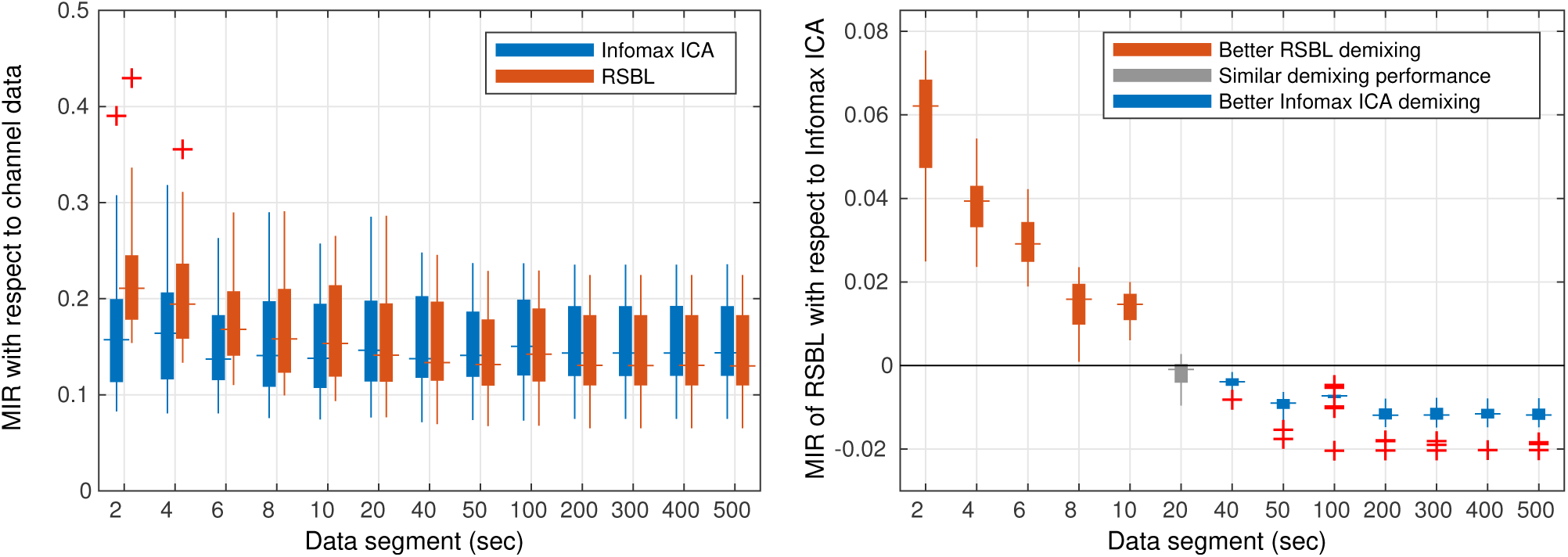
Source separation performance. **Left:** Box plot of MIR with respect to channel data computed on blocks of various sizes. **Right:** Box plot of MIR of RSBL with respect to Infomax ICA for the same data blocks shown on the left. On each box, the central mark indicates the median, and the bottom and top edges indicate the 25th and 75th percentiles respectively. The whiskers extend to the most extreme data points not considered outliers, and the outliers are plotted individually using the + symbol. On the right, the distributions with entire positive (orange) or negative (blue) values indicate a significant source crosstalk reduction in favor of the RSBL or Infomax algorithms respectively.

This result suggests that RSBL can resolve transient sources that may be active for short periods of time. This is not surprising because in RSBL we optimally adjust the resolution of the unmixing matrix (matrix **K**_*k*_ in Eq. (18)) on a millisecond time-scale, which is possible because of all the structure built into our model (see Fig. 3). Infomax (and most ICA algorithms) on the other hand, requires larger data blocks to learn a global factorization of mixing matrix and source activations of reasonable quality. Moreover, Fig 10 suggests that the estimation of a global ICA model is suitable for identifying components that remain stationary over the whole experiment, but otherwise, it is suboptimal for capturing transient dynamics such as those important for BCI applications.

### 3.7. MoBI example: study of heading computation during full-body rotations

We finalize this research with an application of the RSBL algorithm to MoBI data. MoBI experiments are notoriously difficult to analyze due to the amount of motion-induced artifacts as well as the presence of transient and stationary brain dynamics of variable duration across trials. Here, we try to replicate the main findings of a study that looked into the dynamics of the retrosplenial cortex (RSC) supporting heading computation during full-body rotations (Gramann et al., 2018).

Heading computation is key for successful spatial orientation in humans and other animals. The registration of ongoing changes in the environment, perceived through an egocentric first-person perspective has to be integrated with allocentric, viewer-independent spatial information to allow complex navigation behaviors. The RSC provides the neural mechanisms to integrate egocentric and allocentric spatial information by providing an allocentric reference direction that contains the subject’s current heading relative to the environment (Byrne et al., 2007). Single-cell recordings in freely behaving animals have shown that the RSC is also implicated in heading computation (Sharp et al., 2001). And although there is fMRI evidence that points to the same conclusion in humans that navigate in a virtual environment (Baumann and Mattingley, 2010), verifying this hypothesis in more naturalistic settings has remained elusive.

Recently, Gramann et al. (2018) used EEG synchronized to motion capture recordings combined with virtual reality (VR) to investigate the role of the RSC in heading computation of actively moving humans. Data were recorded from 19 participants using 157 active electrodes sampled at 1000 Hz and band-pass filtered from 0.016 Hz to 500 Hz using a BrainAmp Move System (Brain Products, Gilching, Germany). 129 electrodes were placed equidistant on the scalp and 28 were placed around the neck using a custom neckband. In that study, data from physically rotating participants were contrasted with rotations based on visual flow. In the physical rotation condition, participants wore a Vive HTC head-mounted display (HTC Vive; 2 × 1080 × 1200 resolution, 90 Hz refresh rate, 110° field of view). They were placed in a sparse VR environment devoid of any landmark information facing an orienting beacon at the beginning of each trial. The beacon was then replaced by a sphere that started rotating around them to the left or the right at a fixed distance with two different, randomly selected, velocity profiles on each trial. Participants were instructed to rotate on the spot to follow the sphere and keep it in the center of their visual field. The sphere movement was completed at an eccentricity randomly selected between 30° and 150° relative to the initial heading. When the sphere stopped, they had to rotate back and press a controller button to indicate when they believed to have reached their initial heading orientation. After the button press, the beacon would reappear and participants had to rotate to face the beacon and to start the next trial. In the joystick rotation condition, participants stood in front of a large TV screen (1.5 m viewing distance, HD resolution, 60 Hz refresh rate, 40″ diagonal size) controlling a gaming joystick to rotate in the same VR environment with an otherwise identical trial structure.

Using an ICA/dipole fitting approach, the data was analyzed with a focus on oscillatory activity of ICs located in or near the RSC. ICs were clustered using repetitive k-means clustering optimized to the RSC as the region of interest. Four subjects without an IC in the RSC were excluded from the analysis (21% of all participants). Subsequently, the wavelet (Morlet) time-frequency decomposition was computed for each IC in the RSC cluster for the rotation periods. The spectral baseline was defined as the 200 msec period before stimulus onset and subtracted from each time-frequency decomposition. To account for different trial durations, single-trial time-frequency maps were linearly time-warped with respect to the presentation of the stimulus and rotation onset and offset to create time-warped event-related spectral perturbations (ERSPs). Using this approach, the data from the RSC cluster in the joystick rotation condition replicated previous studies using desktop navigation protocols and comparable data analysis approaches (Gramann et al., 2010; Chiu et al., 2012; Lin et al., 2015, 2018), exhibiting 1) a theta burst between stimulus onset and movement onset and 2) alpha and beta desynchronization during the rotation. The physical rotation, however, had drastically different properties: no clear theta burst was present before movement onset, and only minor desynchronization in higher beta bands, but synchronization in the alpha and low beta bands after movement onset and delta and theta bands during the rotation (see Fig 11 **A-B**).

**Figure 11:**
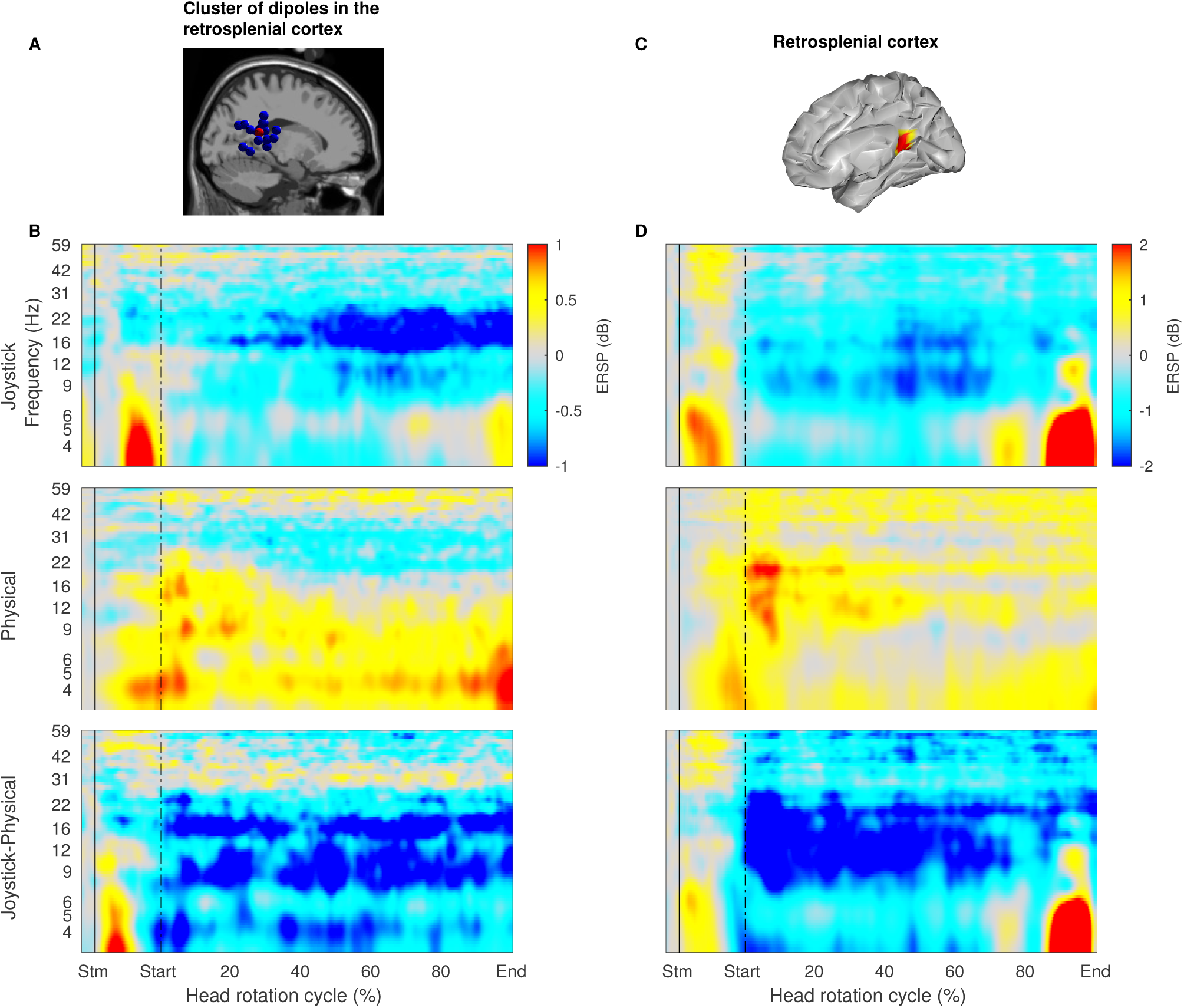
Event-related spectral perturbations (ERSPs) in the RSC. Panels **A** and **B** are adapted from Gramann et al. (2018). **A**: Cluster of IC equivalent current dipoles in or near the RSC. **B**: ICA derived ERSPs of the joystick and physical rotation conditions and their difference. **C**: Location of the RSC in the cortical surface of our template. **D**: RSBL derived ERSPs of the joystick and physical rotation conditions and their difference. The x-axes at the bottom of panels **B** and **D** are annotated with the stimulus onset (Stm), movement onset (Start), percentage of the head rotation cycle, and movement offset (End).

Here, we used the RSBL algorithm to re-analyze the data. To this end, we further down-sampled the data to 250 Hz, removed the neck channels, applied a 0.5 Hz high-pass forward and backward FIR filter, and subtracted the common average reference. We co-registered each subject-specific 129-channels montage with the head surface of the “Colin27” template and computed a lead field matrix and artifact dictionary for each individualized head model. Then we ran the RSBL algorithm for each condition and computed the ERSPs of the centroid source activity averaged within the RSC. The computation of ERSPs was identical to the previous one, only the IC activity of the RSC cluster was replaced by RSBL-resolved RSC source activity of all subjects.

Fig 11 **C-D** shows the RSBL group ERSP for the joystick and physical rotation conditions as well as their difference. The top panel shows in red the location of the RSC in our template brain. Despite the differences between the two methodologies, our results largely replicate those in (Gramann et al., 2018) displayed in panels **A-B**. A few differences between the two results are worth mentioning though. We point out that the differences in ERSP scales exhibited in panels **B** and **D** may be explained by different scales of the sources obtained by ICA and RSBL. Also, we note that the low-frequency power increase towards the end of the head rotation cycle in panel **D** Joystick condition can be explained by artifacts improperly removed near the end of a few trials. It should be emphasized that, unlike the approach used by Gramann et al. (2018), ours has the advantage of using data from all subjects without any pre-cleaning steps in the time, channel, or trial domains, except for the inherent cleaning capabilities of the RSBL algorithm. To increase the robustness to residual artifacts, Fig 8 **D** suggests that a future research direction could explore the use of the log evidence yield by RSBL to automatically downplay the influence of artifactual trials into post hoc statistical summaries.

## 4. Conclusions

In this paper, we have extended the Sparse Bayesian Learning (SBL) framework previously proposed for instantaneous electrophysiological source imaging (Wipf and Nagarajan, 2009) in two ways. First, we augmented the standard generative model of the EEG with a dictionary of artifact scalp projections obtained empirically. In our model, we captured EOG, EMG, and single-channel spike artifacts. Second, we introduced a temporal dynamic constraint and spatial group sparsity constraints based on an anatomical atlas to parameterize the source prior. This parameterization encourages sparsity in the number of active cortical regions, which has the desired property of inducing the segregation of the cortical electrical activity into a few maximally independent components with known anatomical support, while minimally overlapping artifact-related activity. We used these elements to develop the recursive SBL (RSBL) inverse filtering algorithm. Under the proposed framework, dissimilar problems such as data cleaning, source separation, and imaging can be understood and solved in a principled manner using a single algorithm. Furthermore, we used our framework to point out the connections between distributed source imaging and Independent Component Analysis (ICA), two of the most popular approaches for EEG analysis that are often perceived to be at odds with one another.

We used simulated data to show that the RSBL filter can successfully recover the temporal and spatial profile of cortical and artifact sources, even in extremely noisy conditions. We used real data from two independent studies to further test the proposed algorithm. On real data we showed that RSBL: 1) can yield singletrial source estimates of error-related potentials that are in agreement with the experimental literature, 2) can significantly reduce EOG artifacts, 3) unlike the popular Artifact Subspace Removal algorithm, it can reduce artifacts without significantly distorting epochs of clean data, and 4) outperforms Infomax ICA for source separation on short blocks of data, thereby showing potential for tracking non-stationary cortical dynamics. Furthermore, we analyzed MoBI data with RSBL, and we were able to replicate the main finding of a study that investigated the dynamics of the retrosplenial cortex (RSC) supporting heading computation during full-body rotations.

The ability to estimate the time series of EEG sources that correspond to known anatomical locations accounting for the influence of artifacts without user intervention, as well as its online adaptation, makes the RSBL algorithm appealing for established ERP paradigms as well as MoBI. We believe that the proposed algorithm can help to solve basic research questions employing EEG as the functional imaging modality, and at the same time constitute a biologically-grounded signal processing tool that can be useful to translational efforts.

## 5. Acknowledgements

We thank Jason Palmer for sharing his code for computing mutual information reduction. We also thank Rosalyn Moran and Martin Seeber for their valuable comments on an earlier version of this paper. This research was supported by NIMH training fellowships in Cognitive Neuroscience T32MH020002 and Biological Psychiatry and Neuroscience T32MH18399 (AO), UC San Diego Chancellor’s Research Excellence Scholarship (JM, AO), and UC San Diego School of Medicine start-up funds (JM). The RSBL algorithm is copyrighted for commercial use (UC San Diego Copyright #SD2019-810) and free for research and educational purposes.

## Footnotes

1 https://github.com/aojeda/dsi

2 http://bnci-horizon-2020.eu/database/data-sets

## References

Anderson, B.D.O., Moore, J.B., 2012. Optimal Filtering. Dover Publications. reprint of edition.

Baillet, S., Mosher, J.C., Leahy, R.M., 2001. Electromagnetic brain mapping. IEEE Signal Processing Magazine 18, 14–30.

Baumann, O., Mattingley, J.B., 2010. Medial Parietal Cortex Encodes Perceived Heading Direction in Humans. Journal of Neuroscience 30, 12897–901.

Bell, A.J., Sejnowski, T.J., 1995. An information-maximization approach to blind separation and blind deconvolution. Neural Computation 7, 1129–1159.

Bigdely-Shamlo, N., Kreutz-Delgado, K., Kothe, C., Makeig, S., 2013a. EyeCatch: Data-mining over half a million EEG independent components to construct a fully-automated eye-component detector, in: 2013 35th Annual International Conference of the IEEE Engineering in Medicine and Biology Society (EMBC), IEEE. pp. 5845–5848.

Bigdely-Shamlo, N., Mullen, T., Kothe, C., Su, K.M., Robbins, K.A., 2015. The PREP pipeline: standardized preprocessing for large-scale EEG analysis. Frontiers in Neuroinformatics.

Bigdely-Shamlo, N., Mullen, T., Kreutz-Delgado, K., Makeig, S., 2013b. Measure projection analysis: A probabilistic approach to EEG source comparison and multi-subject inference. NeuroImage.

Biscay, R.J., Bosch-Bayard, J.F., Pascual-Marqui, R.D., 2018. Unmixing EEG Inverse solutions based on brain segmentation. Frontiers in Neuroscience.

Böl, M., Weikert, R., Weichert, C., 2011. A coupled electromechanical model for the excitation-dependent contraction of skeletal muscle. Journal of the Mechanical Behavior of Biomedical Materials.

Breakspear, M., 2017. Dynamic models of large-scale brain activity. Nature neuroscience.

Brunner, C., Birbaumer, N., Blankertz, B., Guger, C., Kübler, A., Mattia, D., Millán, J.d.R., Miralles, F., Nijholt, A., Opisso, E., Ramsey, N., Salomon, P., Müller-Putz, G.R., 2015. BNCI Horizon 2020: towards a roadmap for the BCI community. Brain-Computer Interfaces 2, 1–10.

Buzzell, G.A., Richards, J.E., White, L.K., Barker, T.V., Pine, D.S., Fox, N.A., 2017. Development of the error-monitoring system from ages 9âĂŞ35: Unique insight provided by MRI-constrained source localization of EEG. NeuroImage 157, 13–26.

Byrne, P., Becker, S., Burgess, N., 2007. Remembering the past and imagining the future: A neural model of spatial memory and imagery. Psychological Review.

Casella, G., 1985. An introduction to empirical Bayes data analysis. The American Statistician 39, 83–87.

Chang, C.Y., Hsu, S.H., Pion-Tonachini, L., Jung, T.P., 2018. Evaluation of Artifact Subspace Reconstruction for Automatic EEG Artifact Removal, in: 2018 40th Annual International Conference of the IEEE Engineering in Medicine and Biology Society (EMBC), IEEE. pp. 1242–1245.

Chavarriaga, R., del R. Millán, J., 2010. Learning From EEG Error-Related Potentials in Noninvasive Brain-Computer Interfaces. IEEE Transactions on Neural Systems and Rehabilitation Engineering 18, 381–388.

Cheung, B.L.P., Riedner, B.A., Tononi, G., Van Veen, B.D., 2010. Estimation of cortical connectivity from EEG using state-space models. IEEE Transactions on Biomedical Engineering.

Chiu, T.C., Gramann, K., Ko, L.W., Duann, J.R., Jung, T.P., Lin, C.T., 2012. Alpha modulation in parietal and retrosplenial cortex correlates with navigation performance. Psychophysiology.

Cichocki, A., Amari, S.i., 2002. Adaptive Blind Signal and Image Processing.John Wiley & Sons, Ltd, Chichester, UK.

Comon, P., 1994. Independent component analysis, A new concept? Signal Processing 36, 287–314.

Cotter, S., Rao, B.D., Kreutz-Delgado, K., 2005. Sparse solutions to linear inverse problems with multiple measurement vectors. IEEE Transactions on Signal Processing 53, 2477–2488.

Cuspineda, E., Machado, C., Virues, T., Martínez-Montes, E., Ojeda, A., Valdés, P.A., Bosch, J., Valdes, L., 2009. Source Analysis of Alpha Rhythm Reactivity Using LORETA Imaging with 64-Channel EEG and Individual MRI. Clinical EEG and Neuroscience 40, 150–156.

Dale, A.M., Sereno, M.I., 1993. Improved Localizadon of Cortical Activity by Combining EEG and MEG with MRI Cortical Surface Reconstruction: A Linear Approach. Journal of Cognitive Neuroscience 5, 162–176.

Darvas, F., Ermer, J.J., Mosher, J.C., Leahy, R.M., 2006. Generic head models for atlas-based EEG source analysis. Human Brain Mapping 27, 129–143.

Daunizeau, J., Kiebel, S.J., Friston, K.J., 2009. Dynamic causal modelling of distributed electromagnetic responses. NeuroImage 47, 590–601.

Delorme, A., Mullen, T., Kothe, C., Akalin Acar, Z., Bigdely-Shamlo, N., Vankov, A., Makeig, S., 2011. EEGLAB, SIFT, NFT, BCILAB, and ERICA: New tools for advanced EEG processing. Computational Intelligence and Neuroscience 2011.

Delorme, A., Palmer, J., Onton, J., Oostenveld, R., Makeig, S., 2012. Independent EEG sources are dipolar. PLoS ONE.

Desikan, R.S., Ségonne, F., Fischl, B., Quinn, B.T., Dickerson, B.C., Blacker, D., Buckner, R.L., Dale, A.M., Maguire, R.P., Hyman, B.T., Albert, M.S., Killiany, R.J., 2006. An automated labeling system for subdividing the human cerebral cortex on MRI scans into gyral based regions of interest. NeuroImage 31, 968–980.

Friston, K., Harrison, L., Daunizeau, J., Kiebel, S., Phillips, C., Trujillo-Barreto, N., Henson, R., Flandin, G., Mattout, J., 2008. Multiple sparse priors for the M/EEG inverse problem. NeuroImage 39, 1104–1120.

Fujiwara, Y., Yamashita, O., Kawawaki, D., Doya, K., Kawato, M., Toyama, K., Sato, M.a., 2009. A hierarchical Bayesian method to resolve an inverse problem of MEG contaminated with eye movement artifacts. NeuroImage 45, 393–409.

Fukushima, M., Yamashita, O., Kanemura, A., Ishii, S., Kawato, M., Sato, M.A., 2012. A state-space modeling approach for localization of focal current sources from MEG. IEEE Transactions on Biomedical Engineering 59, 1561–1571.

Fukushima, M., Yamashita, O., Knösche, T.R., Sato, M.a., 2015. MEG source reconstruction based on identification of directed source interactions on whole-brain anatomical networks. NeuroImage 105, 408–427.

Galka, A., Yamashita, O., Ozaki, T., Biscay, R., Valdés-Sosa, P., 2004. A solution to the dynamical inverse problem of EEG generation using spatiotemporal Kalman filtering. NeuroImage 23, 435–453.

Gehring, W.J., Liu, Y., Orr, J.M., Carp, J., 2012. The Error-Related Negativity (ERN/Ne), in: The Oxford Handbook of Event-Related Potential Components.

Giraldo, E., den Dekker, a.J., Castellanos-Dominguez, G., 2010. Estimation of dynamic neural activity using a Kalman filter approach based on physiological models. Conference proceedings : … Annual International Conference of the IEEE Engineering in Medicine and Biology Society. IEEE Engineering in Medicine and Biology Society. Conference 2010, 2914–7.

Gorodnitsky, I.F., Rao, B.D., 1997. Sparse signal reconstruction from limited data using FOCUSS: A re-weighted minimum norm algorithm. IEEE Transactions on Signal Processing 45, 600–616.

Gramann, K., Ferris, D.P., Gwin, J., Makeig, S., 2014. Imaging natural cognition in action. International Journal of Psychophysiology 91, 22–29.

Gramann, K., Hohlefeld, F.U., Gehrke, L., Klug, M., 2018. Heading computation in the human retrosplenial complex during full-body rotation. bioRxiv.

Gramann, K., Onton, J., Riccobon, D., Mueller, H.J., Bardins, S., Makeig, S., 2010. Human brain dynamics accompanying use of egocentric and allocentric reference frames during navigation. Journal of Cognitive Neuroscience.

Gramfort, A., Papadopoulo, T., Olivi, E., Clerc, M., 2010. Open-MEEG: opensource software for quasistatic bioelectromagnetics. Biomedical engineering online 9, 45.

Gramfort, a., Strohmeier, D., Haueisen, J., Hämäläinen, M.S., Kowalski, M., 2013. Time-frequency mixed-norm estimates: Sparse M/EEG imaging with non-stationary source activations. NeuroImage 70, 410–22.

Hallez, H., Vanrumste, B., Grech, R., Muscat, J., De Clercq, W., Vergult, A., D’Asseler, Y., Camilleri, K.P., Fabri, S.G., Van Huffel, S., Lemahieu, I., 2007. Review on solving the forward problem in EEG source analysis. Journal of Neuro-Engineering and Rehabilitation 4, 46.

Hämäläinen, M.S., Ilmoniemi, R.J., 1994. Interpreting magnetic fields of the brain: minimum norm estimates. Medical & Biological Engineering & Computing 32, 35–42.

Haufe, S., Tomioka, R., Dickhaus, T., Sannelli, C., Blankertz, B., Nolte, G., Müller, K.R., 2011. Large-scale EEG/MEG source localization with spatial flexibility. NeuroImage 54, 851–859.

Henson, R.N., Wakeman, D.G., Litvak, V., Friston, K.J., 2011. A Parametric Empirical Bayesian Framework for the EEG/MEG Inverse Problem: Generative Models for Multi-Subject and Multi-Modal Integration. Frontiers in Human Neuroscience 5.

Hsu, S.H., Mullen, T.R., Jung, T.P., Cauwenberghs, G., 2016. Real-Time Adaptive EEG Source Separation Using Online Recursive Independent Component Analysis. IEEE Transactions on Neural Systems and Rehabilitation Engineering.

Huang, M.X., Dale, A.M., Song, T., Halgren, E., Harrington, D.L., Podgorny, I., Canive, J.M., Lewis, S., Lee, R.R., 2006. Vector-based spatial-temporal minimum L1-norm solution for MEG. NeuroImage.

Huang, Y., Parra, L.C., Haufe, S., 2016. The New York HeadâĂŤA precise standardized volume conductor model for EEG source localization and tES targeting. NeuroImage.

Islam, M.K., Rastegarnia, A., Yang, Z., 2016. Methods for artifact detection and removal from scalp EEG: A review. Neurophysiologie Clinique/Clinical Neurophysiology.

Janani, A.S., Grummett, T.S., Lewis, T.W., Fitzgibbon, S.P., Whitham, E.M., DelosAngeles, D., Bakhshayesh, H., Willoughby, J.O., Pope, K.J., 2017. Evaluation of a minimum-norm based beamforming technique, sLORETA, for reducing tonic muscle contamination of EEG at sensor level. Journal of Neuroscience Methods 288, 17–28.

Jung, T.P., Makeig, S., Humphries, C., Lee, T.W., Mckeown, M.J., Iragui, V., Sejnowski, T.J., 2000. Removing electroencephalographic artifacts by blind source separation. Psychophysiology

Jungnickel, E., Gramann, K., 2016. Mobile Brain/Body Imaging (MoBI) of Physical Interaction with Dynamically Moving Objects. Frontiers in Human Neuroscience 10.

Kalman, R.E., 1960. A New Approach to Linear Filtering and Prediction Problems. Journal of Basic Engineering.

Khambhati, A.N., Sizemore, A.E., Betzel, R.F., Bassett, D.S., 2018. Modeling and interpreting mesoscale network dynamics. NeuroImage 180, 337–349.

Kilicarslan, A., Grossman, R.G., Contreras-Vidal, J.L., 2016. A robust adaptive denoising framework for real-time artifact removal in scalp EEG measurements. Journal of Neural Engineering 13, 026013.

Lamus, C., Hämäläinen, M.S., Temereanca, S., Brown, E.N., Purdon, P.L., 2012. A spatiotemporal dynamic distributed solution to the MEG inverse problem. NeuroImage.

Le, Q.V., Karpenko, A., Ngiam, J., Ng, A., 2011. ICA with reconstruction cost for efficient overcomplete feature learning, in: Shawe-Taylor, J., Zemel, R., Bartlett, P., Pereira, F., Weinberger, K. (Eds.), Advances in Neural Information Processing Systems 24, Curran Associates, Inc.. pp. 1017–1025.

Lewicki, M.S., Sejnowski, T.J., 1998. Learning nonlinear overcomplete representations for efficient coding. Advances in Neural Information Processing Systems 10 (NIPS’97), 556–562.

Lin, C.T., Chiu, T.C., Gramann, K., 2015. EEG correlates of spatial orientation in the human retrosplenial complex. NeuroImage.

Lin, C.T., Chiu, T.C., Wang, Y.K., Chuang, C.H., Gramann, K., 2018. Granger causal connectivity dissociates navigation networks that subserve allocentric and egocentric path integration. Brain Research.

Long, C.J., Purdon, P.L., Temereanca, S., Desai, N.U., Hämäläinen, M.S., Brown, E.N., 2011. State-space solutions to the dynamic magnetoencephalography inverse problem using high performance computing. Annals of Applied Statistics.

MacKay, D.J.C., 1992. Bayesian Interpolation. Neural Computation 4, 415–447.

MacKay, D.J.C., 1996. Hyperparameters: Optimize, or integrate out? Maximum entropy and bayesian methods, 43–59.

MacKay, D.J.C., 2008a. Independent Component Analysis and Latent Variable Modelling, in: Information Theory, Inference, and Learning Algorithms. Cambridge University Press, pp. 437–440.

MacKay, D.J.C., 2008b. Information Theory, Inference, and Learning Algorithms. Cambridge University Press.

Makeig, S., Gramann, K., Jung, T.P.P., Sejnowski, T.J., Poizner, H., 2009. Linking brain, mind and behavior. International Journal of Psychophysiology 73, 95–100.

Makeig, S., Jung, T.P., Bell, A.J., Ghahremani, D., Sejnowski, T.J., 1997. Blind separation of auditory event-related brain responses into independent components. Proceedings of the National Academy of Sciences of the United States of America.

Makeig, S., Onton, J., 2011. ERP Features and EEG Dynamics. Oxford University Press.

Mannan, M.M.N., Kamran, M.A., Jeong, M.Y., 2018. Identification and Removal of Physiological Artifacts From Electroencephalogram Signals: A Review. IEEE Access 6, 30630–30652.

Martínez-Vargas, J.D., Grisales-Franco, F.M., Castellanos-Dominguez, G., 2015. Estimation of M/EEG Non-stationary Brain Activity Using Spatio-temporal Sparse Constraints, in: Artificial Computation in Biology and Medicine. Springer International Publishing, pp. 429–438.

Mcdowell, K., Lin, C.T., Oie, K.S., Jung, T.P., Gordon, S., Whitaker, K.W., Li, S.Y., Lu, S.W., Hairston, W.D., 2013. Real-world neuroimaging technologies. IEEE Access.

Mehta, R.K., Parasuraman, R., 2013. Neuroergonomics: a review of applications to physical and cognitive work. Frontiers in Human Neuroscience.

Michel, C.M., Murray, M.M., 2012. Towards the utilization of EEG as a brain imaging tool. NeuroImage 61, 371–385.

Mishra, J., Anguera, J.A., Gazzaley, A., 2016. Video Games for Neuro-Cognitive Optimization.

Mishra, J., Gazzaley, A., 2014. Closed-Loop Rehabilitation of Age-Related Cognitive Disorders. Seminars in Neurology 34, 584–590.

Morris, C.N., 1983. Parametric empirical Bayes inference: theory and applications. Journal of the American Statistical Association 78, 47–55.

Mullen, T.R., Kothe, C.A.E., Chi, Y.M., Ojeda, A., Kerth, T., Makeig, S., Jung, T.P., Cauwenberghs, G., 2015. Real-time neuroimaging and cognitive monitoring using wearable dry EEG. IEEE Transactions on Biomedical Engineering 62, 2553–2567.

Neal, R.M., 1996. Bayesian Learning for Neural Networks. volume 118 of Lecture Notes in Statistics. Springer New York, New York, NY.

Nunez, P.L., Srinivasan, R., 2006. Electric Fields of the Brain. Oxford University Press.

Ojeda, A., Kreutz-Delgado, K., Mullen, T., 2018. Fast and robust Block-Sparse Bayesian learning for EEG source imaging. NeuroImage 174, 449–462.

Olier, I., Trujillo-Barreto, N.J., El-Deredy, W., 2013. A switching multi-scale dynamical network model of EEG/MEG. NeuroImage 83, 262–287.

Onton, J., Westerfield, M., Townsend, J., Makeig, S., 2006. Imaging human EEG dynamics using independent component analysis. Neuroscience & Biobehavioral Reviews 30, 808–822.

Owen, J.P., Wipf, D.P., Attias, H.T., Sekihara, K., Nagarajan, S.S., 2012. Performance evaluation of the Champagne source reconstruction algorithm on simulated and real M/EEG data. NeuroImage 60, 305–323.

Ozaki, T., 2012. Time Series Modeling of Neuroscience Data. CRC Press.

Palmer, J., Kreutz-Delgado, K., Makeig, S., 2011. AMICA: An Adaptive Mixture of Independent Component Analyzers with Shared Components. San Diego, CA: Technical report, Swartz Center for Computational Neuroscience.

Pascual-Marqui, R.D., Esslen, M., Kochi, K., Lehmann, D., 2002. Functional imaging with low-resolution brain electromagnetic tomography (LORETA): a review. Methods and findings in experimental and clinical pharmacology 24 Suppl C, 91–5.

Penny, W., Friston, K., Ashburner, J., Kiebel, S., Nichols, T., 2007. Statistical Parametric Mapping. Elsevier.

Pion-Tonachini, L., Makeig, S., Kreutz-Delgado, K., 2017. Crowd labeling latent Dirichlet allocation. Knowledge and Information Systems 53, 749–765.

Radüntz, T., Scouten, J., Hochmuth, O., Meffert, B., 2017. Automated EEG artifact elimination by applying machine learning algorithms to ICA-based features. Journal of Neural Engineering.

Sharp, P.E., Blair, H.T., Cho, J., 2001. The anatomical and computational basis of the rat head-direction cell signal. Trends in Neurosciences 24, 289–294.

Lopes da Silva, F., 2013. EEG and MEG: Relevance to Neuroscience. Neuron 80, 1112–1128.

Tamburro, G., Fiedler, P., Stone, D., Haueisen, J., Comani, S., 2018. A new ICA-based fingerprint method for the automatic removal of physiological artifacts from EEG recordings. PeerJ 6, e4380.

Tipping, M.E., 2001. Sparse Bayesian Learning and the Relevance Vector Machine. Journal of Machine Learning Research 1, 211–245.

Treder, M.S., Bahramisharif, A., Schmidt, N.M., Van Gerven, M.A., Blankertz, B., 2011. Brain-computer interfacing using modulations of alpha activity induced by covert shifts of attention. Journal of NeuroEngineering and Rehabilitation.

Trujillo-Barreto, N.J., Aubert-Vázquez, E., Penny, W.D., 2008. Bayesian M/EEG source reconstruction with spatio-temporal priors. NeuroImage 39, 318–335.

Trujillo-Barreto, N.J., Aubert-Vázquez, E., Valdés-Sosa, P.A., 2004. Bayesian model averaging in EEG/MEG imaging. NeuroImage 21, 1300–1319.

Valdés-Hernández, P.A., von Ellenrieder, N., Ojeda-Gonzalez, A., Kochen, S., Alemán-Gómez, Y., Muravchik, C., Valdés-Sosa, P.A., 2009. Approximate average head models for EEG source imaging. Journal of Neuroscience Methods 185, 125–132.

Valdes-Sosa, P.A., Sanchez-Bornot, J.M., Sotero, R.C., Iturria-Medina, Y., Aleman-Gomez, Y., Bosch-Bayard, J., Carbonell, F., Ozaki, T., 2009. Model driven EEG/fMRI fusion of brain oscillations.

Valdés-Sosa, P.A., Vega-Hernández, M., Sánchez-Bornot, J.M., Martínez-Montes, E., Bobes, M.A., 2009. EEG source imaging with spatio-temporal tomographic nonnegative independent component analysis. Human Brain Mapping 30, 1898–1910.

Van Der Maaten, L., Hinton, G., 2008. Visualizing Data using t-SNE. Journal of Machine Learning Research.

Van Veen, B.D., van Drongelen, W., Yuchtman, M., Suzuki, A., 1997. Localization of brain electrical activity via linearly constrained minimum variance spatial filtering. IEEE Transactions on Biomedical Engineering.

Wagner, J., Makeig, S., Gola, M., Neuper, C., Muller-Putz, G., 2016. Distinct Band Oscillatory Networks Subserving Motor and Cognitive Control during Gait Adaptation. Journal of Neuroscience 36, 2212–2226.

Ward, T., Bernier, R., Mukerji, C., Perszyk, D., McPartland, J.C., Johnson, E., Faja, S., Nevers, M., Frazier, T., Howlin, P., Savage, S., Zane, T., Lanner, T., Myers, M., VanBergeijk, E., Huestis, S., Bauminger-Zviely, N., Doehring, P., Voorst, G., Macy, K., Kwon, J.M., McNulty, E., Chapman, S.M., Crowley, M.J., Bean, A., Hyman, S., Scahill, L.D., Wing, L., Catania, A.C., Thorne, J., Kini, U., Moyle, M., Plowgian, C., Happé, F., Case-Smith, J., Schmitt, L., Lewis, M., Weiss, M.J., Gaag, J.V.d.R., Hurst, H., Bonazinga, L., Paul, D.R., Happé, F., Doyle, C.A., McDougle, C.J., Perri, K.S., Moran, M., Stigler, K., McDougle, C.J., Hyman, S., St. John, M., Hessl, D., Schneider, A., Katon, J., Hurst, H., Bauminger-Zviely, N., Hadjikhani, N., Rinehart, N., Enticott, P., Bradshaw, J., Palmieri, M., LaRue, R., Molteni, J., LaRue, R., Palmieri, M., Prelock, P., Müller, R.A., LaRue, R., Egan, S., Hendricks, D., Holman, K.C., DâĂZEramo, K., Faja, S., Perszyk, D., 2013. Feedback-Related Negativity, in: Encyclopedia of Autism Spectrum Disorders. Springer New York, New York, NY, pp. 1256–1257.

Winkler, I., Haufe, S., Tangermann, M., 2011. Automatic Classification of Artifactual ICA-Components for Artifact Removal in EEG Signals. Behavioral and Brain Functions 7, 30.

Wipf, D., Nagarajan, S., 2009. A unified Bayesian framework for MEG/EEG source imaging. NeuroImage 44, 947–966.

Wipf, D.P., Nagarajan, S.S., 2008. A New View of Automatic Relevance Determination, in: Advances in Neural Information Processing Systems, pp. 1625–1632.

Yamashita, O., Galka, A., Ozaki, T., Biscay, R., Valdes-sosa, P., 2004. Recursive Penalized Least Squares Solution for Dynamical Inverse Problems of EEG Generation. Human Brain Mapping 235, 221–235.

Yang, Y., Aminoff, E.M., Tarr, M.J., Kass, R.E., 2016. A state-space model of cross-region dynamic connectivity in MEG/EEG. Advances In Neural Information Processing Systems.

Zhang, Z., Rao, B.D., 2013. Extension of SBL Algorithms for the Recovery of Block Sparse Signals With Intra-Block Correlation. IEEE Transactions on Signal Processing 61, 2009–2015.

